# Understanding species-specific and conserved RNA-protein interactions *in vivo* and *in vitro*

**DOI:** 10.1101/2024.01.29.577729

**Authors:** Sarah E. Harris, Maria S. Alexis, Gilbert Giri, Francisco F. Cavazos, Jernej Murn, Maria M. Aleman, Christopher B. Burge, Daniel Dominguez

## Abstract

While evolution is often considered from a DNA- and protein-centric view, RNA-based regulation can also impact gene expression and protein sequences. Here we examined interspecies differences in RNA-protein interactions using the conserved neuronal RNA binding protein, Unkempt (UNK) as model. We find that roughly half of mRNAs bound in human are also bound in mouse. Unexpectedly, even when transcript-level binding was conserved across species differential motif usage was prevalent. To understand the biochemical basis of UNK-RNA interactions, we reconstituted the human and mouse UNK-RNA interactomes using a high-throughput biochemical assay. We uncover detailed features driving binding, show that *in vivo* patterns are captured *in vitro*, find that highly conserved sites are the strongest bound, and associate binding strength with downstream regulation. Furthermore, subtle sequence differences surrounding motifs are key determinants of species-specific binding. We highlight the complex features driving protein-RNA interactions and how these evolve to confer species-specific regulation.

## Introduction

Species divergence and adaptation rely on a delicate balance of robustness — the ability to withstand mutations without serious deleterious effects on fitness — and evolvability — the susceptibility to developing a novel phenotype^1,2^. Driving this balance are changes in gene expression programs and coding sequences^2–5^. Understanding how changes in *trans* (nucleic acid-binding proteins) and *cis* (nucleic acid sequences) impact gene regulation across species remains an important challenge. While changes in *cis* and *trans* across species are both important, *cis*-regulatory elements change more rapidly than *trans* factor amino acids or binding preferences^6,7^. For example, transcription factors (TFs) and RNA-binding proteins (RBPs) are highly conserved over long evolutionary distances while regions harboring *cis*-regulatory elements that these proteins bind can vary drastically over the same distances^8,9^.

Most previous studies have taken a DNA- and protein-centric view (reviewed by Villar *et al*.^10^ and Mitsis *et al*.^11^); however, RNA regulation influences both expression levels as well as protein coding sequences, resulting in potential widespread effects^12–14^. RBPs constitute a large class of pan-essential regulatory factors^15,16^ that drive RNA regulation, contributing significantly to transcription, splicing, and trans-lation to influence the expression and identity of proteins produced^17–20^. These processes are dictated by the strength of the interaction between RBPs and their RNA targets^21–25^. In the simplest model of RNA regulation, RBPs bind short sequence motifs (3-8 nucleotides) within RNA to influence its regulation^26^. However, these interactions are complex as change, loss, or gain of a single nucleotide within or surrounding a motif can greatly impact binding^27–29^.

RBPs themselves have a striking level of amino acid conservation with many RNA-binding domains (RBDs) remaining nearly identical, even over hundreds of millions of years^30^. Generally, RBPs tend to be more conserved than their DNA-binding counterparts, transcription factors^9^. Paradoxically, RNA processing events regulated by RBPs, such as alternative splicing and translation, have been found to be more species-specific and to evolve more rapidly than gene expression programs (*i*.*e*., tissue-specific expression across species)^31–34^. How often are regulatory elements that control gene expression and RNA processing conserved across species? If binding has changed, what are the mechanisms underlying that change? Previous studies on TF binding to regulatory elements have addressed this in a number of species^6,7,35–38^ (and reviewed by Villar *et al*.^10^). Multiple studies — in-cluding one employing chromatin immunoprecipitation and sequencing (ChIP-Seq) across five vertebrates — have found that although TFs are highly conserved, *cis*-regulatory elements evolve rapidly and primarily dictate TF binding profiles^6^. More specifically, TF binding profiles (*i*.*e*., bound genes) demonstrate less than 40% conservation between human and mouse, even though the individual TFs studied are nearly identical (>95% identity) at the amino acid level and have identical or near-identical binding preferences^7^. When a human chromosome is placed in a mouse context, TF binding predominantly follows the human binding patterns rather than that of the mouse^35^, indicating that binding pattern changes are primarily *cis*-directed. Of course, these forms of interactome evolution are partially dependent on evolutionary time and the individual TFs being assessed^39^.

Few similar studies of species-specific RNA-protein interactions have been conducted. But some emerging themes parallel the similarities to that of TF-DNA interactions. For example, previous work has examined the conservation of the Pumilio and FBF (Puf) superfamily of proteins and their interactomes^40–43^. Puf3 exhibits highly similar RNA binding specificities across fungal species^42^; however, Puf3 targets change significantly between *S. cerevisiae* and *N. crassa*^43^. More strikingly, targets bound by Puf3 in one species are bound by a different RBP — Puf4/5 — in another^43^, highlighting the complex nature of RNA binding site evolution and the interplay between *cis* and *trans*. Within species, single nucleotide polymorphisms (SNPs) have been shown to impact RBP-RNA interactions. A comprehensive analysis of RBP-RNA interaction studies in two cell types identified over a thousand cases of allele-specific RBP-RNA interactions, some of which were validated biochemically and had functional impact on RNA regulation^44^. The complex paths in which RNA regulation evolves have been previously reviewed^45^, but much work is still needed to understand how the underlying driving forces, namely RBP-RNA interactions, drive changes in regulation.

To understand species-specific RNA binding, we used available individual-nucleotide resolution crosslinking and immunoprecipitation (iCLIP) data^27^ from a neuronal RBP, unkempt (UNK), in human and mouse. UNK regulates neuronal morphology, is a negative regulator of translation, and associates with polysomes^27,46,47^. We identified species-specific and shared UNK binding sites and found that 45% of UNK transcript binding was conserved across species. Importantly, while binding of transcripts was conserved, the individual motifs that were bound were far less conserved, often switching between species. We reconstituted the *in vivo* UNK-RNA interactomes of human and mouse *in vitro* to understand the driving forces underlying species-specific binding and regulation. We found that while motif turnover is an important mediator of species-specific binding, contextual sequence and structural features in which motifs are embedded are of comparable importance and contribute to binding site turnover. We extended our studies across 100 vertebrates to understand how sequence changes over longer time scales affect binding and found striking correlations between evolutionary distances, individual binding site conservation, and strength of UNK binding. This work deepens our understanding of *cis*-regulatory evolution and highlights the complex nature of evolving RNA binding sites.

## Results

### Conserved and Species-Specific *in vivo* Binding Patterns

We undertook an RNA-centric view and sought to determine how RNA binding sites change or are conserved across species. We focused on the conserved neuronal RBP, unkempt (UNK) for the following reasons: i) UNK has a well-defined RNA-binding motif supported by structural studies^46^; ii) UNK is 95% conserved between human and mouse with only one amino acid difference within the RNA-binding zinc finger domains^48–50^; iii) Murn and coworkers demonstrated that even the sea sponge (*Amphimedon queenslandica*) UNK paralog functionally rescues knockdown of UNK in human cell lines^46^ even though these species only share 53% similarity at the protein level and 80% similarity within the RBDs^48–50^. Thus, this level of functional conservation provides an opportunity to study changes in UNK binding sites across species primarily driven by changes in RNA sequence rather than in the protein’s binding properties. We used UNK iCLIP data in human and mouse neuronal cells and tissue, respectively^27^, to identify species-specific and conserved UNK binding sites (**Fig. S1A**). Comparing binding sites across species at the transcript level, we observe that 45% of transcripts are bound in both species (**Fig. 1A**-Venn diagram; p=6e-94, hypergeometric test). As iCLIP allows for individual nucleotide level determination of binding sites, we further investigated where on each transcript UNK was bound. UNK binding sites require a UAG core motif, which has been identified both *in vitro* and *in vivo*^26,27^. In instances where transcript level binding was conserved between human and mouse, roughly half of binding was observed at aligned (homologous) motifs across species. In cases where binding sites within transcripts changed across species, motif loss only accounted for a minority of these changes. That is, in many cases both human and mouse preserved a UAG motif in the same location, yet binding was often identified elsewhere on the transcript (**Fig. 1A**-top pie chart). Likewise, when comparing motif differences in transcripts only bound in human or mouse, motifs were preserved across species in over 70% of orthologous regions (**Fig. 1A**-bottom pie charts) yet binding was differential. Thus, while UNK protein is highly conserved, engagement of UNK with specific UAG motifs often varies across species.

**Figure 1.**
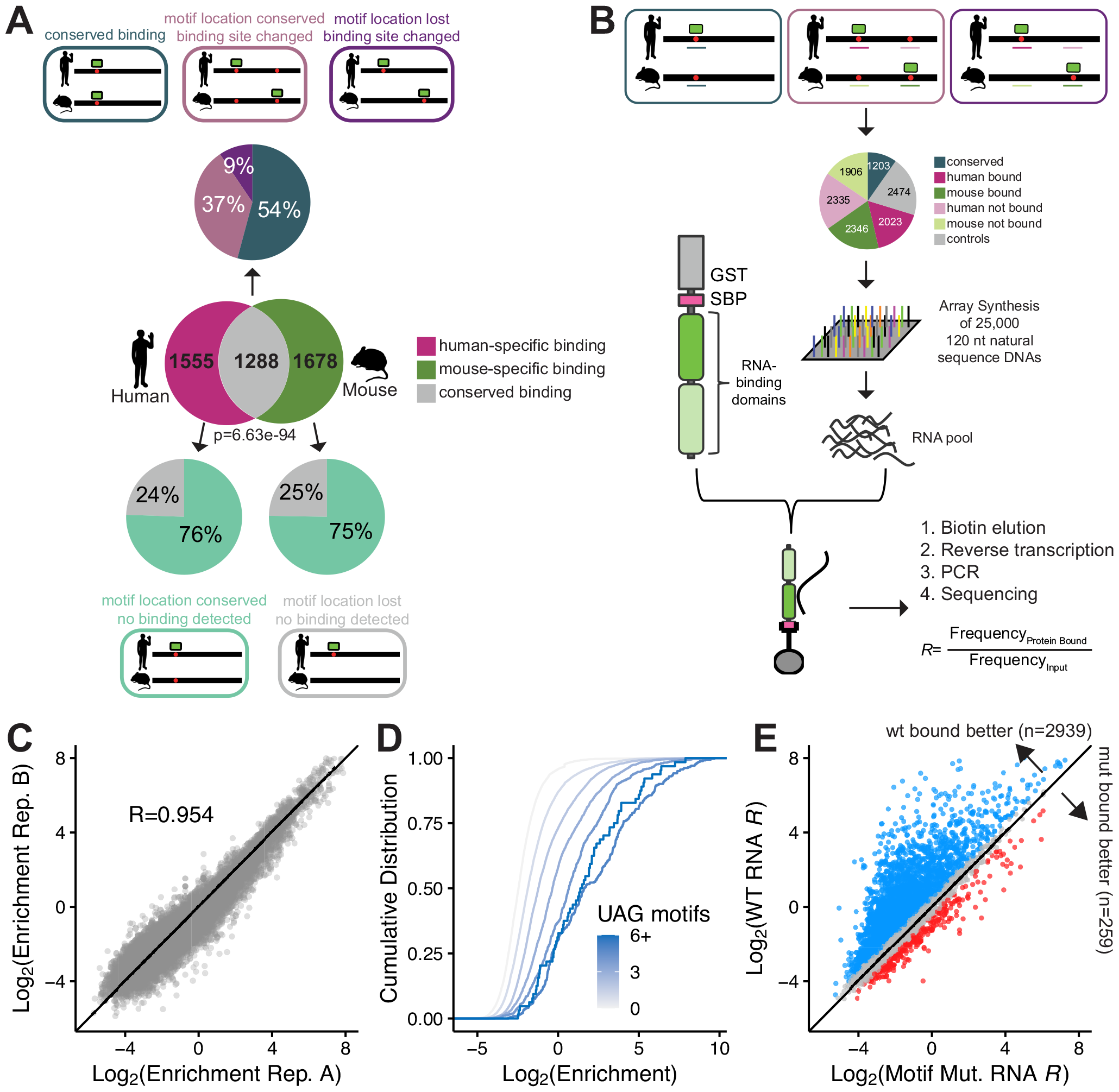
Design and validation of natural sequence RNA bind-n-seq (nsRBNS). A) (Venn diagram) Transcript level conservation of iCLIP UNK hits between human neuronal cells (SH-SY5Y) and mouse brain tissue. Significance determined via hypergeometric test. (Pie charts) Motif level conservation of iCLIP UNK hits between human neuronal cells (SH-SY5Y) and mouse brain tissue. B) Design of natural sequence oligo pool and layout of nsRBNS. C) Correlation plot of two experimental UNK nsRBNS replicates. Pearson’s correlation coefficient included. D) Cumulative distribution function of log_2_ enrichment of all oligos separated by UAG motif content. E) Scatter plot of log_2_ enrichment of wildtype (Y-axis) versus motif mutant (X-axis) oligos. Log_2_ change in enrichment (wt-mut) was calculated for each sequence pair: > 0.5 defined as bound better in wt (blue), < -0.5 defined as bound better in mut (red), 0 ± 0.5 defined as similar binding (grey).

### Understanding the UNK-RNA Interactome *in vitro* at Massive Scale

While iCLIP is a powerful technique that allows for the derivation of nucleotide-level binding sites, several experimental factors, including RNA crosslinking efficiencies and biases across RBPs, cell types, and tissues, complicate its interpretation. Another important consideration for understanding binding site conservation across species via iCLIP is that false negative rates of CLIP experiments are largely unknown^51^. Finally, the strength of RBP-RNA interactions found in CLIP-based experiments have a limited dynamic range. That is, binding affinity and occupancy are not easily determined, and binding is often interpreted as binary when a continuum of occupancy levels likely occur *in vivo*. To mitigate CLIP biases and understand binding differences across species due to the intrinsic properties of the RNA-protein interactions, we sought to reconstitute the human and mouse UNK-RNA interactomes *in vitro*.

We derived UNK binding sites from iCLIP data in one-to-one orthologous human and mouse genes (see **Methods**) and designed 12,287 natural RNA sequences, each 120 nucleotides long. Contained within this “pool” were UNK binding sites identified via iCLIP in human and mouse neuronal cells and tissue, respectively, as well as orthologous regions whether or not they displayed evidence of binding in cells (**Fig. 1B**). Sequences were designed such that UAGs identified via iCLIP were located in the center of each oligo whenever possible (**Methods**). Non-bound control regions were also selected and matched for UAG content (**Fig. 1B**). Additionally, 11,967 mutated oligos were also included and are discussed below.

An array of these natural sequence DNA oligos was synthesized and underwent *in vitro* transcription to generate an RNA pool. To determine how UNK protein binds these 25,000 sequences, we performed natural sequence RNA Bind-n-Seq^26,52^ (nsRBNS), a quantitative large-scale *in vitro* binding assay (**Fig. 1B**). Briefly, the RNA pool of natural sequences was incubated with recombinant protein, protein-RNA complexes were immobilized on magnetic beads, washed, and RNA was isolated. RNA sequencing was used to quantify the abundance of each RNA bound to UNK as well as the abundance of each RNA in the input RNA pool. These experiments yield binding enrichments (*R* values) for each oligo which are defined as the frequency of a given oligo bound to UNK vs the frequency of that oligo in the input RNA (**Methods**). Greater *R* values indicate a higher degree of binding. Previous work has demonstrated that nsRBNS correlates well with *in vivo* binding and regulation^53,54^. This approach enabled us to test the binding of nearly 25 thousand sequences in tandem, with a wide range of *in vivo* binding properties.

UNK nsRBNS experiments were performed in duplicate and at different protein concentrations with robust cross-replicate correlation (**Fig. 1C**; R=0.954, Pearson’s correlation). We first asked whether nsRBNS is capable of capturing binding differences based on previously derived UNK motifs. UNK is known to bind a primary core UAG motif with secondary U/A-rich motifs^26,27^. Presence of more than one UNK motif within an RNA has been shown to enhance binding^26^, driven by engagement with the tandem zinc fingers of UNK^46^. Within our pool, we observe that binding enrichment increased with increasing UAG count (**Fig. 1D**), consistent with previous studies^26^. Similar but slightly more modest increases in binding occurred with increasing counts of UUU and UUA (**Fig. S1B**,**C**).

To demonstrate that the core UAG motif is important for binding, we included central motif mutants. For these sequences, if there was a central UAG motif present within the 120 nt region, it was mutated to CCG to assess whether binding is reduced (**Methods**). As expected, and as reported previously^27^, mutating the central UAG motif is enough to drastically diminish binding (**Fig. 1E**). This observation was further validated via an *in vitro* qPCR-based binding assay for one gene, *GART*. We observed that mutation of the central UAG motif to a CCG drastically diminished binding (**Fig. S1D**; p=0.004, one-sided Wilcox test). Finally, given that UNK is known to bind single-stranded RNA^26^, we computed the base-pairing probability of the central 10 nt region harboring binding sites using a thermodynamic RNA folding algorithm^55^. As the mean base-pairing probability of this central region decreased (e.g., more structure occluding the region) enrichment values decreased (**Fig. S1E**). These data confirm nsRBNS as a replicable *in vitro* assay, capable of measuring binding differences based on sequence features for 25,000 sequences in parallel.

### Recapitulation of *in vivo* Binding Patterns and Regulation

We next tested whether *in vivo* binding patterns could be recapitulated *in vitro*. For our *in vitro* analysis, we defined three classes of binding sites: “control” where no evidence of binding was detected via iCLIP in either species; “bound,” where binding was detected via iCLIP; and “orthologous not bound,” where sites were bound in one species and not bound in the other (**Fig. 2A**, diagram). As UNK has been shown to bind primarily within the coding sequences (CDS) and secondarily within 3’ untranslated regions (UTRs)^27^, we assessed binding patterns individually for these regions. Within CDS binding sites, we found that “orthologous not bound” oligos had similar enrichments as “control” oligos whereas “bound” oligos were significantly more enriched (**Fig. 2B**, p<2.2e-16, KS test). In UTRs, “bound” oligos were again the most enriched, but in this case “orthologous not bound” sites in UTRs had greater enrichments than controls (**Fig. 2C**). In fact, UTR sites overall had better enrichments than CDS, perhaps due to UTRs being generally enriched for U- and A-rich 3mers that are bound by UNK (**Fig. S2A**). Thus, nsRBNS captures binding features derived from in vivo iCLIP.

**Figure 2.**
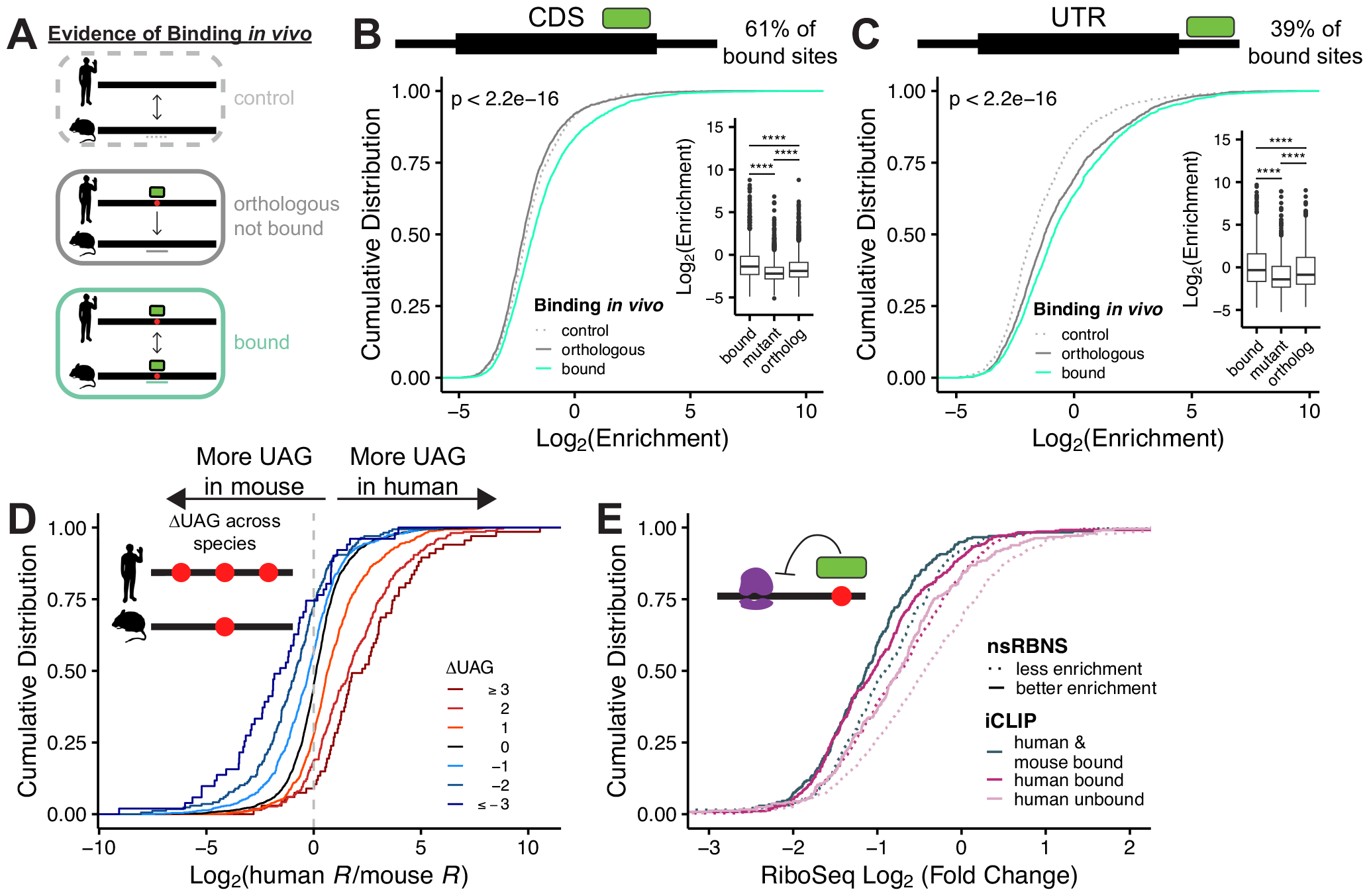
Analysis of species-specific binding patterns. A) Schematic of “control,” “orthologous,” and “bound” oligo classes used for species-specific transcript-level binding analysis. B-C) Cumulative distribution function of log_2_ enrichment of all iCLIP hits: control (light grey; dotted), orthologous (dark grey), and bound (teal) of B) CDS and C) UTR oligos. Significance of bound vs. orthologous was determined via KS test. Insets show boxplot of *in vitro* binding patterns for “bound,” “motif mutant,” and “orthologous” oligos. Significance was determined via two-sided Wilcox test. D) Cumulative distribution function of log_2_ fold enrichment change of *in vivo* bound over *in vivo* not bound oligos separated by ΔUAG content. E) Cumulative distribution function of RiboSeq log_2_ fold change separated via iCLIP detection. nsRBNS enrichment cutoffs defined as “less enrichment” <1 and “better enrichment” >1.

nsRBNS enrichment values span several orders of magnitude, driven by differences in affinity and avidity (**Fig. 1D**). Although occurring in a far more complex environment, binding in cells likely also occurs on a spectrum driven by affinity, though more difficult to capture experimentally. To compare *in vivo* to *in vitro* patterns, we asked what proportion of species-specific binding observed *in vivo* could be captured *in vitro*. We measured how often a species-specific site was better bound than its non-bound ortholog and found that 60% of binding sites mirrored the *in vivo* trend (**Fig. 2B**,**C**-inset & **Fig. S2B**,**C**). However, the degree to which species-specific binding was recapitulated *in vitro* ranged from no difference to greater than 100-fold difference between the bound and unbound orthologous site. To better understand these patterns, we turned to *in vivo*-bound sites where we also mutated the UAG motif (see above **Fig. 1E**). We reasoned that because UAG drives binding, these mutants would be representative of minimal binding. Indeed, in 80% of cases UAG mutation diminished binding (**Fig. 2B**,**C**-inset & **Fig. S2B**,**C**). In aggregate, we found that “orthologous not bound” sites had an intermediate enrichment, that is, not as weakly bound as UAG mutants but significantly less bound *in vitro* than the “bound” category. Consistent with these findings and what is known about UNK-RNA interactions, the difference in UAG content between human and mouse orthologous sites had a large impact on differential binding (**Fig. 2D**), with gain of UAG enhancing binding and loss decreasing binding. The same was true of the known secondary motifs UUU (**Fig. S2D**) and UUA (**Fig. S2E**). Additionally, the greater the difference in percent identity between the 120 nt human and mouse binding sites, the greater the absolute difference in enrichments across species (**Fig. S2F**,**G**). These data highlight that *in vivo* binding patterns can be recapitulated *in vitro*. However, we note that some *in vivo* differences are not captured *in vitro*, likely reflecting a combination of the cellular environment and limitations of *in vivo* (CLIP) and *in vitro* (nsRBNS) assays.

### Binding Strength and *in vivo* Regulation

To determine whether these *in vitro* binding patterns also correspond with *in vivo* regulatory patterns, we examined ribosome profiling data upon UNK induction^47^. UNK is a translational repressor^27^, thus UNK-regulated RNAs are predicted to have decreased translation as previously shown^27^. Genes with peaks identified via iCLIP^27^ in both human and mouse were more translationally repressed than genes with peaks only identified in human or genes lacking UNK peaks (**Fig. 2E**). These data highlight that conserved targets display stronger regulation, consistent with what has been observed for microRNAs and splicing factors^56–59^. We next asked whether the strength of binding from *in vitro* (nsRBNS) also had an impact on regulation. Greater *in vitro* binding enrichments were associated with increased translational suppression for transcripts that were common to mouse and human or to human alone (**Fig. 2E**). These data support a direct relationship between interactions measured *in vitro* and binding and regulation assessed *in vivo*.

### Species-Specific Binding Site Patterns

As noted above, iCLIP analysis revealed that while binding can be conserved at the transcript level, specific binding locations often change (**Fig. 3A**). For example, within the *GGPS1* transcript, we observe two species-specific binding sites (**Fig. 3B**). *In vivo*, this transcript is bound in both human and mouse, but different binding sites are used (located 122 nt apart in the alignment) even though the UAG motifs are conserved (**Fig. 3B**, bottom). In our *in vitro* nsRBNS assay, we see enrichments that mirror these *in vivo* patterns (**Fig. 3B**, top) prompting us to use this biochemical data to examine binding site preferences.

**Figure 3.**
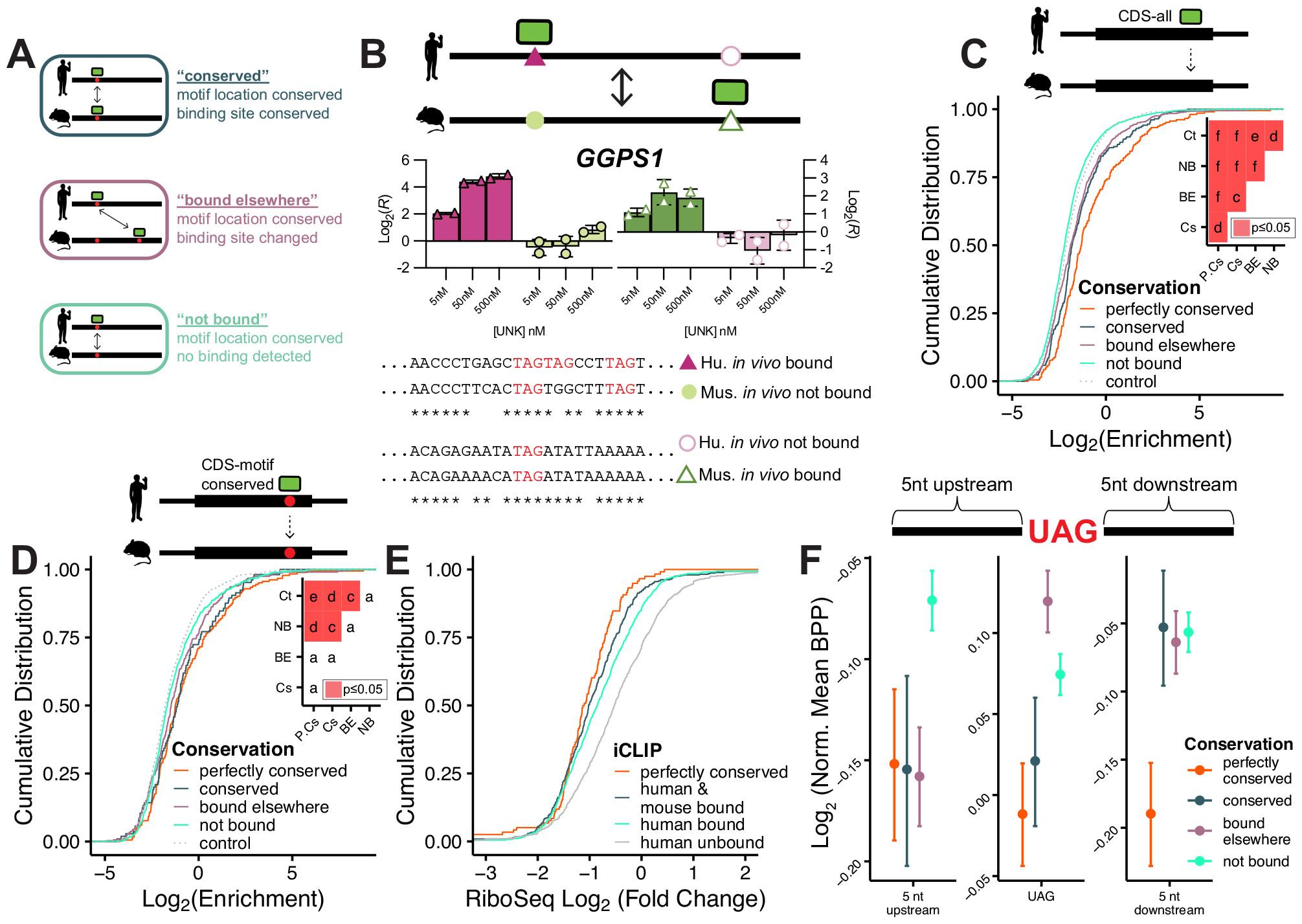
Analysis of species-specific syntenic motif level binding patterns. A) Definition of “conserved,” “bound elsewhere,” and “not bound” oligo classes used for species-specific transcript regional binding analysis. B) Conservation and binding of *GGPS1* orthologous pairs. (left) Log_2_ enrichment values from nsRBNS for human bound (purple triangle), mouse not bound (light green circle), mouse bound (green open triangle), and human not bound (light purple open circle). (right) Alignment of human bound (purple triangle) to mouse not bound (light green circle) and mouse bound (green open triangle) to human not bound (purple open circle). Note: full oligos were used for alignment but only central region is shown. C-D) Cumulative distribution function of log_2_ enrichment of control (light grey; dotted), not bound (teal), bound elsewhere (purple), conserved (blue), and perfectly conserved (orange) C) all CDS and D) motif conserved CDS oligos. Insets show significance values for all comparisons via KS test and corrected for multiple comparisons via the BH procedure. Red denotes significant (p<=0.05). Values are as follows: a (ns), c (p<=0.05), d (p<=0.01), e (p<=0.001), f (p<=0.0001). E) Cumulative distribution function of RiboSeq log_2_ fold change separated via iCLIP detection and sequence conservation. F) Log_2_ fold change of mean base pair probability of the central region of “perfectly conserved,” “conserved,” “bound elsewhere,” and “not bound,” oligos normalized to UAG-containing CDS controls (see **Methods**). Error bars show standard error of the mean.

To determine how binding location changes across species in an *in vitro* context, we included four additional classes of oligos within our pool: “conserved,” where both the motif and binding location were maintained across species; “bound elsewhere,” where transcript binding was conserved, yet there was still differential motif usage across species (even when a motif was preserved across species); “not bound,” where the motif was maintained yet binding was not detected in the orthologous species (**Fig. 3A**); and “perfectly conserved,” the subset of “conserved” oligos with identical sequences between human and mouse. In aggregate, the degree of conserved binding *in vivo* correlated with *in vitro* enrichments. Least enriched were the “not bound” category followed by “bound elsewhere,” then “conserved,” and most enriched were “perfectly conserved” sites (**Fig. 3C**,**D & Fig. S3A**,**B**). Surprisingly, *in vitro* binding followed *in vivo* binding even when only regions with UAG motifs in both human and mouse were considered (**Fig. 3D & Fig. S3B**). Similar trends were observed in CDS and UTR regions (CDS in **Fig. 3C**,**D**; UTR in **Fig. S3A**,**B**). These data demonstrate that factors beyond the core motif impact RBP-RNA interactions, as our data shows that UNK can switch UAG motif usage between species. The fact that these preferences can be captured *in vitro* indicates that *cis* sequence changes surrounding the motifs are an important driver of binding.

Of note, binding sites perfectly conserved (100% identity) between human and mouse were among the strongest bound. In fact, of these regions we found that fewer than 3% were bound in only one species *in vivo*, indicating that a high degree of conservation within larger sequence regions is associated with conserved binding. To associate this degree of conservation with *in vivo* regulation, we again turned to ribosome profiling after UNK induction and found that transcripts with perfectly conserved binding sites were more translationally suppressed than other bound transcripts (**Fig. 3E**).

We hypothesized that these high-affinity binding sites may be more accessible (*i*.*e*., have reduced levels of RNA secondary structure). To this end, we aligned sequences by their central UAG, performed in silico folding^55^, and compared each of the above categories. Indeed, perfectly conserved binding sites were the most accessible (with lower base pair probabilities (BPP)) at and downstream of the motif (**Fig. 3F**). Consistent with the preferences of these and many other RBPs for single-stranded RNA^26^, accessibility appears to drive evolutionary changes in RNA binding. Simply put, conservation of context is a critically important mediator of conserved RNA-protein interactions.

### Binding Site Patterns Across Cell Types Within Species

To compare these binding preferences to intra-species changes, we examined available iCLIP data from HeLa cells overexpressing UNK from the same study^27^. Surprisingly, when looking at transcript-level conservation, we observed that approximately 51% of UNK transcripts were bound in both cell types (**Fig. S3C**; p=2.3e-202, hypergeometric test), similar to that observed in human vs. mouse comparisons. Looking further at the binding site level, only 41% occurred at the same motif (**Fig. S3D**), again similar to the cross-species comparisons. However, when we turned to *in vitro* nsRBNS, sites that were bound in both cell types versus only bound in one had no biochemical difference in binding as enrichments were largely similar (**Fig. S3E**). These data suggest that differing cellular environments (likely including presence of different complements of RBPs) can influence binding locations to a substantial degree.

To compare these patterns more generally to a larger group of RBPs across cell types within humans, we assessed binding of 14 RBPs (with well-defined motifs) from available enhanced CLIP (eCLIP) data in HepG2 and K562 cells^60,61^ (**Fig. S3F**). Although eCLIP differs from iCLIP^61,62^ (reviewed by Hafner *et al*.^63^), we reasoned that both types of experiments should yield similar information. Using these data, we found that RBP binding sites — although variable from RBP to RBP — are well-conserved at the transcript level across cell types with 64% conservation on average for exonic binding and 53% conservation on average for non-exonic binding (e.g., introns) between HepG2 and K562 cells. At the binding site level, approximately 54% of exonic peaks and 41% of non-exonic peaks are bound at the same motif across cell types (**Fig. S3G**). As expected, peaks with well-defined motifs displayed a greater degree of overlap between cell lines (**Fig. S3H**,**I**). These observations are similar to what we observed for UNK between SH-SY5Y versus HeLa cells with iCLIP (**Fig S3C**,**D**). Thus, although limited to a small cohort of RBPs, these data suggest that whereas inter-species differences can be largely influenced by *cis* changes that can be captured biochemically (as discussed above), intra-species differences may be dictated by changing cellular environment across tissues (*i*.*e*., RNA/RBP expression levels, levels of other RBPs, etc.).

### Species-Specific Regional Impacts on Binding

When binding is species-specific, is it possible to identify the sequences that drive binding in one species but not the other? In the simplest scenario this would be a region harboring a UAG motif that is found in only one species. To test whether introduction of sequences from an *in vivo*-bound species to the orthologous region that displayed no *in vivo* binding could restore binding, we designed chimeric mutants. Starting with the unbound mouse sequence, we substituted 10 nucleotide segments of the bound sequence into the unbound sequence to test which parts, if any, of the human sequence could confer binding (**Fig. 4A**).

**Figure 4.**
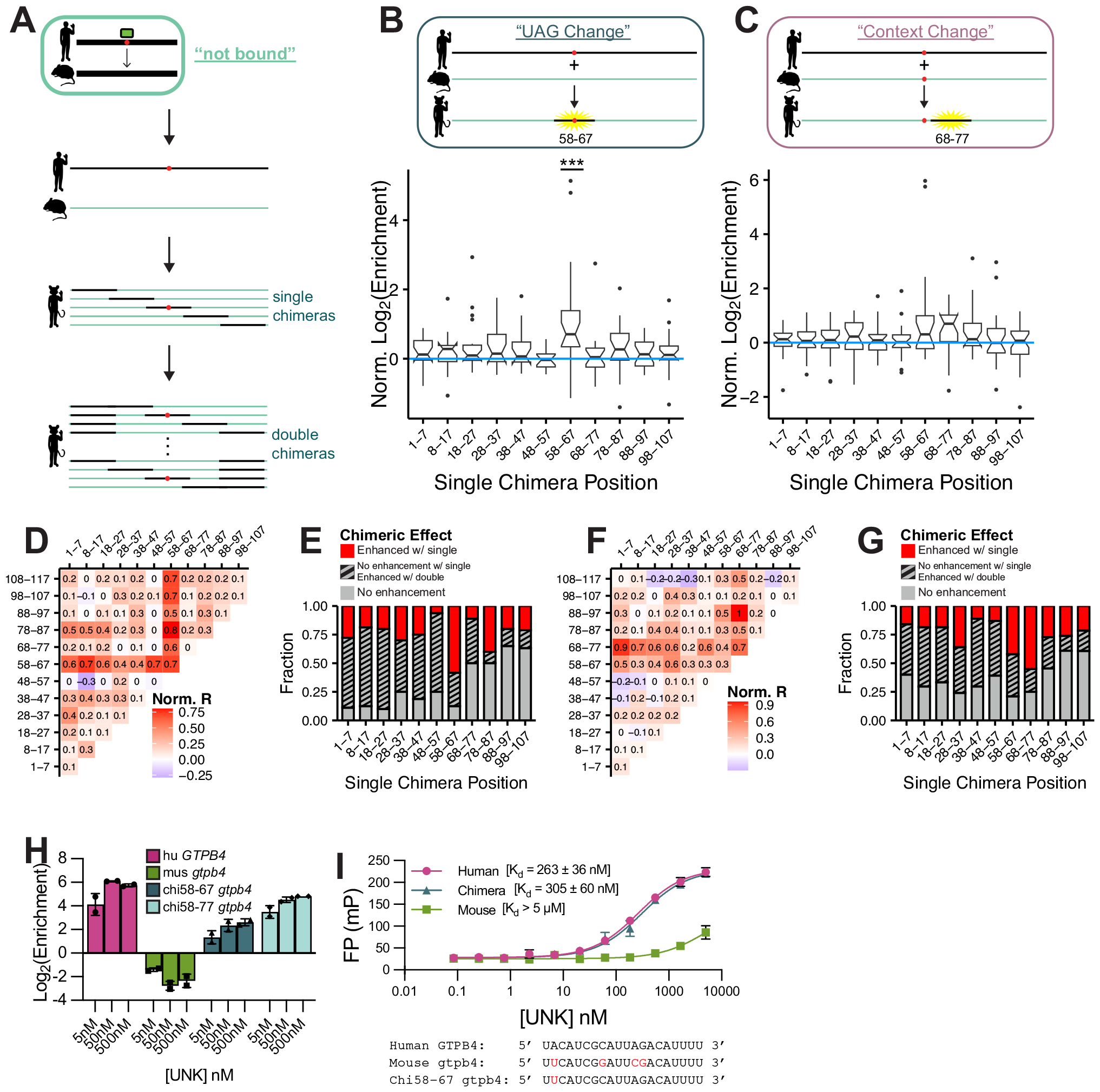
Analysis of regional impacts on binding. A) Design of single and double chimera oligos. B) Design and box and whisker plot of normalized log_2_ enrichment (chimera/wt) for “UAG Change” single chimeras. Significance was determined via paired, one-sided Wilcox test and corrected for multiple comparisons via the BH procedure. Chimerization at positions 58-67 was found to be significant (p<=0.001). C) Design and box and whisker plot of normalized log_2_ enrichment (chimera/wt) for “Context Change” single chimeras. Significance was determined via paired, one-sample Wilcox test and corrected for multiple comparisons via the BH procedure. Following multiple comparison correction, no positions were determined to be significant. D) Heat map of median normalized log_2_ enrichment (chimera/wt) for “UAG Change” single and double chimeras. E) Fraction of “UAG Change” chimeras enhanced with binding after single (red) or double (grey, striped) chimerization. F) Heat map of median normalized log_2_ enrichment (chimera/wt) for “Context Change” single and double chimeras. G) Fraction of “Context Change” chimeras enhanced with binding after single (red) or double (grey, striped) chimerization. H) Log_2_ enrichment values from nsRBNS for human, mouse, single chimera, and double chimera *GTPB4* at 5, 50, and 500 nM UNK. I) Fluorescence polarization binding curves for human *GTPB4*, mouse *gtpb4*, and chi58-67 *gtpb4* RNA oligos incubated with UNK.

Within these chimeric oligos we included two classes: “UAG Change,” where the central UAG was present in the bound sequence but not in the unbound mouse sequence; and “Context Change,” where the UAG was conserved in both. As expected, in a “UAG Change” example, substitution of the central 10 bases which include the UAG motif (58-67) significantly enhanced binding (**Fig. 4B**). Supporting the importance of contextual features, other positions not harboring the central UAG could also confer enhanced binding but no single chimerized position contributed as significantly as position 58-67 which harbored the UAG (**Fig. 4B**).

Of particular interest were the “not bound” cases where a motif was conserved across species yet binding was lost (**Fig. 3A**). In “Context Change” chimeras, we noted a boost in binding upon changing of positions 58-67 (that harbor the central UAG) despite the motif being present in both species, suggesting contextual differences. Importantly, we also found swapping the segment just downstream (68-77) appeared to enhance binding (**Fig. 4C**), though statistical significance was not reached after correcting for multiple tests in this small cohort of binding sites tested (n=22). Enhancing chimeric sequences — mostly downstream of the core motif — tend to be U/A rich (**Fig. S4A**), likely leading to increased avidity and further engagement of UNK’s secondary RBD^46^.

RNA binding is complicated and multi-factorial. Therefore, we hypothesized that chimerization of 10 nucleotides might not be sufficient to recover binding across species and that longer-range effects could be at play. Thus, we also included double chimeras where every possible combination of single chimeras was tested for binding (**Fig. 4A**). As we expected, double chimerization of both “UAG Change” (**Fig. 4D**) and “Context Change” (**Fig. 4F**) cases improved binding in many cases where single substitutions did not. Interestingly, almost all combinations with position 58-67 for “UAG Change” chimeras significantly enhanced binding (**Fig. S4B-D**). Double chimerization enhanced binding for 70% of “UAG Change” chimeras whereas single chimerization only achieved 21% restoration (**Fig. 4E**). Likewise, for “Context Change” double chimerization enhanced binding up to 52% from 24% with single chimeras (**Fig. 4G**). For approximately 50% of double chimeras, double chimerization not only restores binding but also led to binding at the level of the bound human sequence (**Fig. S4E**,**F**).

In one example, UNK bound human *GTPB4* approximately 500-fold better than mouse *Gtpb4*. Single chimerization of the central UAG-containing region (58-67) restored binding 30-fold (**Fig. 4H**), while double chimerization with positions 68-77 improved binding an additional 3.5-fold (**Fig. 4H**). We validated these binding patterns for *GTPB4* via fluorescence polarization (FP) with a 6-FAM-tagged RNA and fit a Kd for human *GTPB4* at 262.6 nM, mouse *Gtbp4* at >5 μM, and chi58-*Gtpb4* at 304.8 nM (**Fig. 4I**). Thus, evolution of UNK-RNA binding involves substantial contributions of both primary motif level and contextual changes.

### Evolutionary Conservation of Binding

To expand our phylogenetic scope beyond human/mouse, we investigated binding patterns across 100 vertebrates. Selecting the top 250 bound human sequences from our original nsRBNS assay, we sought to identify orthologous regions from 100 vertebrates^64,65^, keeping only those where 25 or more species were aligned (**Fig. 5A & S5A**; see **Methods**). Within this set of sequences, we also included total motif mutants for each human oligo where every UAG was mutated to a CCG to define cutoffs for null binding (**Methods**). We performed nsRBNS as described above (**Fig. S5B**). As an initial analysis, we measured the decrease in binding between wild-type regions and total motif mutant regions in human. As expected, we found that wild-type human sequences are better bound across the assay than mutant counterparts (**Fig. S5C**; p=1.1e-15, two-sided, paired Wilcoxon test).

**Figure 5.**
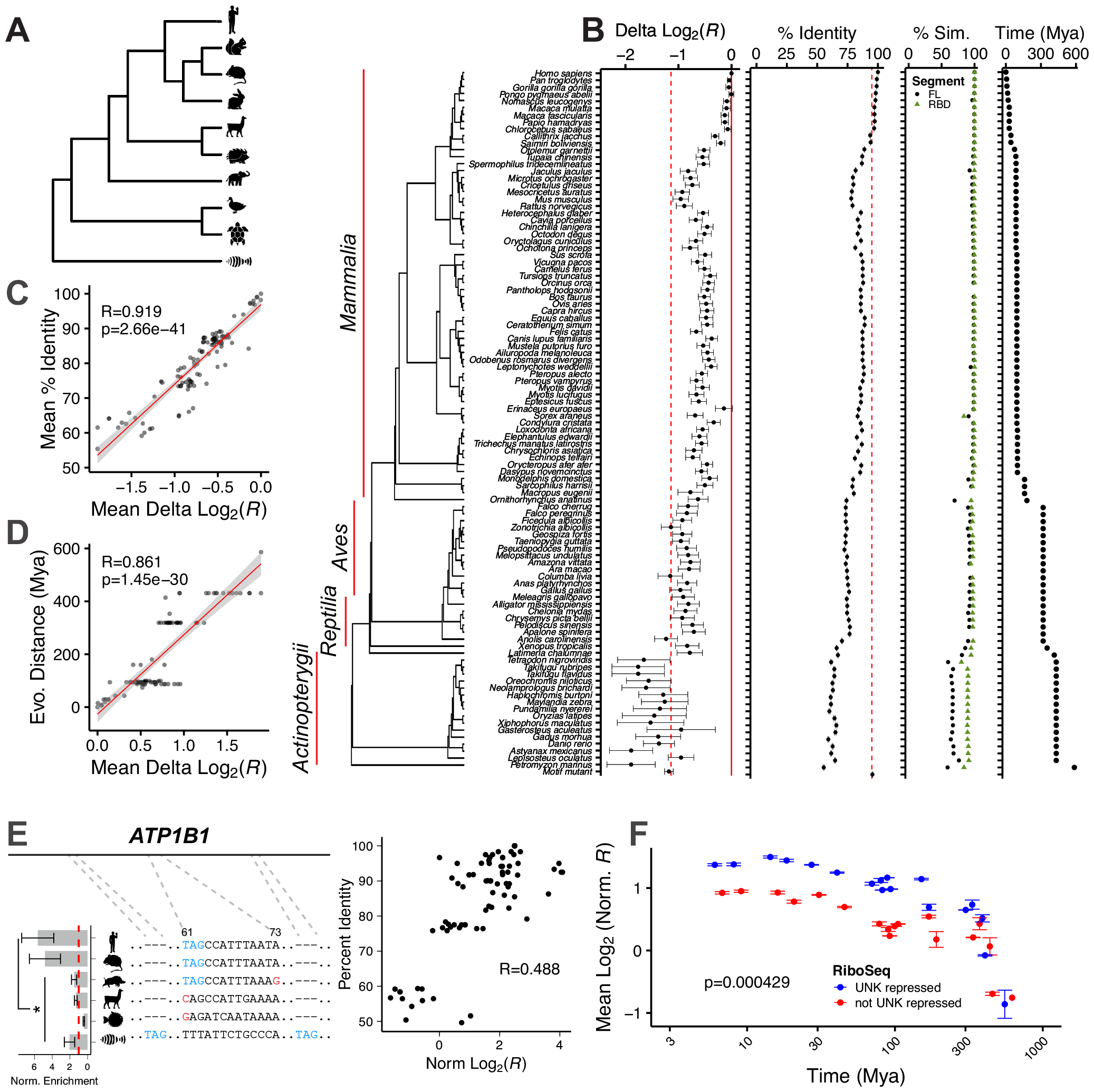
Evolutionary Conservation of Binding. A) Simplified alignment of 100 vertebrate natural sequence alignment. B) Delta log_2_ enrichment, percent RNA sequence identity, percent UNK similarity (full length-grey and RBDs-green), and evolutionary distance in millions of years against 100 vertebrates for the aligned sequences from the top human bound oligos. Error bars show standard error of the mean (SEM). C) Mean percent RNA sequence identity (Y-axis) versus mean delta log_2_ enrichment (X-axis) for each aligned oligo. Pearson’s correlation coefficient included. D) Evolutionary distance in millions of years (Y-axis) versus mean delta log2 enrichment (X-axis) for each aligned oligo. Pearson’s correlation coefficient included. E) (left) Multiple sequence alignment for *ATP1B1* for *Homo sapiens, Mus musculus, Sus scrofa, Vicugna pacos, Tetradon nigroviridis*, and *Danio rerio* with normalized enrichment by species. (right) Percent RNA sequence identity (Y-axis) versus normalized delta log_2_ enrichment (X-axis). Pearson’s correlation coefficient included. F) Scatter plot of log_2_ normalized UNK binding enrichment by evolutionary distance. X-axis plotted on log_10_ scale. Error bars show SEM. Data were separated by regulation as determined via RiboSeq where blue reflects UNK repression of translation [higher than average log_2_ fold change (>-0.9)] and red reflects lack of UNK repression [less than average log2 fold change (<-0.9)]. Significance was determined via KS test.

To understand how binding diverges along the evolutionary timeline, we set the difference between each wild-type region and its UAG-mutant as the dynamic range of max binding to no binding (**Methods**). As we progress to more distant species from human, we observed that binding enrichment decreased (**Fig. 5B**). To understand the driving force behind loss or maintenance of binding, we computed sequence identity between human and all other vertebrates tested for every binding site (**Fig. 5B**, center). Some species and families along the tree have RNA sequence identity more similar to human which is mirrored by an increase in binding enrichment (**Fig. 5B**, center). These changes are reflected with a remarkably high degree of correlation between mean percent identify to human and binding enrichments (**Fig. 5C**; R=0.919, Pearson’s correlation). Similarly, evolutionary distances are also well-correlated with binding (**Fig. 5D**; R=0.861, Pearson’s correlation). A large drop-off in binding with increased variance was observed in fish (class *Actinopterygii*) (**Fig. 5B**, left), suggesting a loss of human UNK compatibility and species-specific RNA-protein interactions. Not surprisingly, however, while the RNA sequences evolve rapidly, with percent identity dropping quickly, UNK protein is highly conserved across species and does not reach 60% similarity (compared to human) until pufferfish (*Tetraodon nigroviridis*; 431 million years divergence), while the RBDs never drop below 70% similarity in vertebrates^48,49^ (**Fig. 5B**, center right).

In one example, *ATP1B1*, the central UAG motif is well conserved through pufferfish; however, binding begins to drop off around Cape golden mole (*Chrysochloris asiatica*). Similar to the trend for all binding sites tested, the percent identity of *ATP1B1* orthologs to human positively correlated with UNK binding (**Fig. 5E & Fig. S5D**,**E**). While this binding dropoff is not mirrored by any apparent changes in UAG content, a subtle shift in downstream sequence may be responsible for this binding difference. This can be observed through wild boar (*Sus scrofa*) *Atp1b1* which still maintains a central UAG motif but has a decrease in A/U content just downstream. Interestingly, alpaca (*Vicugna pacos*) *Atp1b1* binds with similar enrichment to wild boar *Atp1b1* even though it has completely lost its central UAG motif, perhaps indicating that the downstream changes in A/U content in wild boar are as important as loss of the UAG. At further evolutionary distances, zebrafish (*Danio rerio*) has completely lost the UAG that confers binding in human but has gained a UAG motif upstream and downstream which is mirrored through an increase in binding enrichment. Similarly complex binding trajectories are observed for *NFATC3* orthologs (**Fig. S5F-H**). While many primate sequences were perfectly identical, we observed with *PPP2R5C* that even across short evolutionary distances, large motif changes can occur, leading to drastic changes in binding (**Fig. S5I**,**J**).

To examine how binding changes across 100 vertebrates correlates with RNA regulation, we once again turned to ribosome profiling data^47^. As discussed above, we found correlations between UNK binding and evolutionary distance and sequence conservation (**Fig. 5F**). Interestingly, binding sites within mRNAs that UNK translationally suppressed displayed a higher degree of UNK binding conservation across vertebrates (**Fig. 5F**), spanning large evolutionary distances. This effect may be explained by the fact that translational targets were modestly more conserved than those that are not translational targets (**Fig. S5K**).

These data support a model wherein both subtle (often difficult to discern) and large changes greatly influence RNA-protein interactions and therefore RNA regulation (**Fig. 6**). Often adjacent sequences and RNA structure change — sometimes driven by single substitutions — resulted in loss of binding *in vitro*. These subtle sequence differences can seemingly have impacts akin to total motif loss, implicating these differences in loss of *in vivo* binding. Future work will expand these studies beyond UNK, however, given the prototypical nature of UNK-RNA interactions and the high degree of conservation at the protein level, we expect that other RBPs behave similarly.

**Figure 6.**
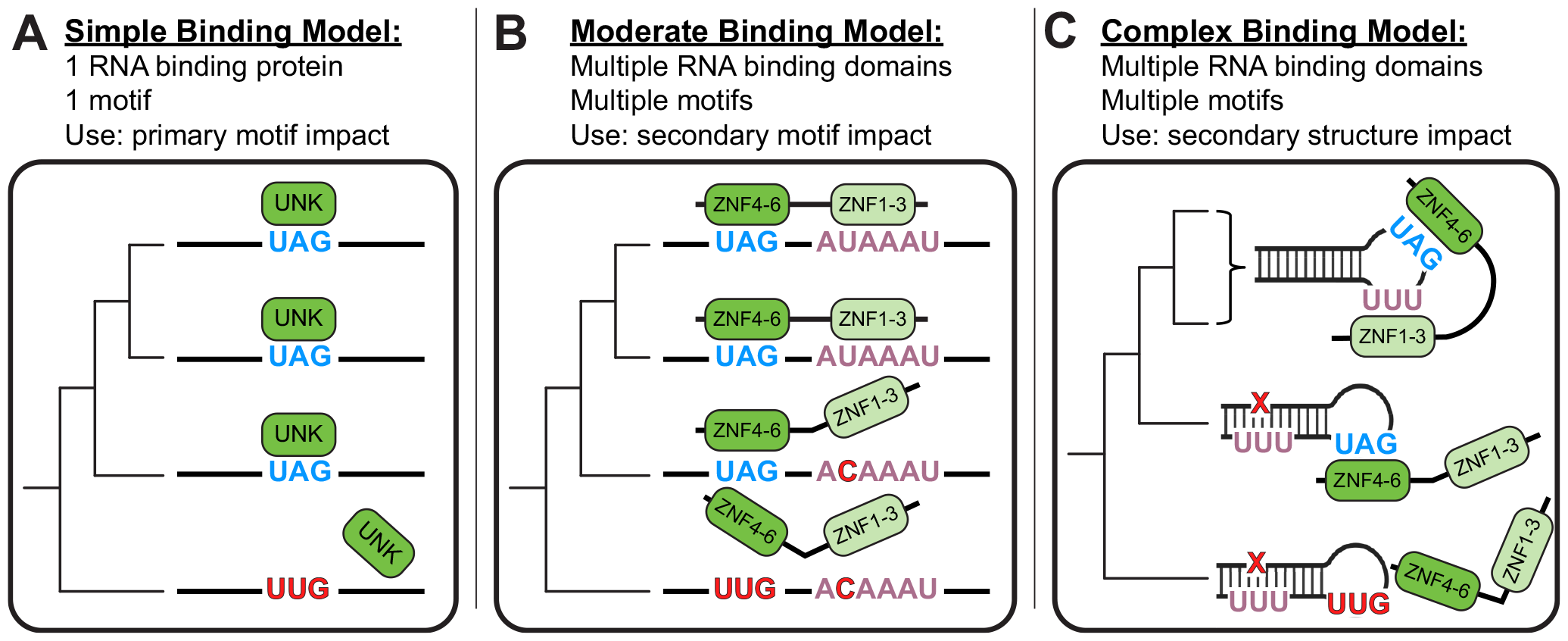
Models of RNA Binding. A) Simple binding model: considers only primary motifs. B) Moderate binding model: considers primary and secondary motifs. C) Complex binding model: considers primary and secondary motifs as well as RNA secondary structure.

### Limitations of This Study

Our approach relies on assessing how well *in vivo* data is reflected using high-throughput *in vitro* approaches. This strategy is well-suited for understanding direct protein-RNA interactions influenced by changes in RNA sequence and local RNA structure; however, it does not include the complex cellular environment including factors that can impact binding *in vivo*: **i)** Individual mRNA and RBP concentrations can vary drastically in cells, across cell types, and across species. The concentration of the RNA, the RBP, other components of mRNPs, and ribosomal interactions (as UNK is primarily involved in translational control^27^) can all be anticipated to affect UNK-RNA interactions and accessibility (see **Supp. Note 1**). **ii)** Our work has been guided by *in vivo* binding data, which — while incredibly informative — have limitations (discussed above) which ultimately prompted us to reconstitute the UNK-RNA interactome *in vitro*. Because these iCLIP experiments partially guided our experimental designs, technical biases present in iCLIP may have impacted our choice of sequences to study (see **Supp. Note 1**). **iii)** Due to difficulties in purifying full-length UNK protein, we have performed our experiments on the RBDs alone, which exhibit tight and specific binding to known UNK motifs (as shown above and previously^26^); additional components of UNK, such as its disordered domain, may impact binding in unexpected ways^47^. **iv)** For practical reasons we have used human UNK protein for these studies. For smaller evolutionary distances such as human versus mouse, this is unlikely to impact binding. However, for larger distances where UNK protein RBDs may be more different, one might expect to see co-evolution of the *cis*-elements with changes in RNA-binding properties of UNK (see **Supp. Note 1**). **v)** RBNS is in nature a zero-sum experiment in that all RNA molecules compete with each other for protein binding. Thus, in pools that contain mostly high affinity RNA targets, some targets will appear enriched and others depleted despite all being high-affinity binders. This effect is most evident in our assays using binding sites derived from 100 vertebrates (**Fig. S5B**; see **Supp. Note 2**). **vi)** The nature of oligo pool design removed sequences with poor alignments. Therefore, the percent identity analysis shown is reflective of only those binding sites that had a minimal level of alignment (**Methods**), while those not having sufficient alignments were excluded. The above should be considered when interpreting enrichments presented in this study.

## Discussion

How differences in RNA sequence impact RNA-protein interactions and downstream regulation remains poorly understood. Until recently, studies focused primarily on TFs and their binding sites; however, recent work has begun to incorporate studies on RNA^31,32,40–45,66^. Here, we examine species-specific and conserved RNA binding using the neuronal RBP, UNK, as a model. We find that roughly 45% of UNK binding sites have been maintained between human and mouse, and far fewer maintain these binding sites down to the motif level (**Fig. 1**). Using a high-throughput *in vitro* assay to test binding, we found that species-specific binding patterns and regulation can be explained to a substantial effect by biochemically measurable RNA-protein interactions (**Fig. 2 & Fig. 3**). Although more limited in scope, binding changes across cell types within species appear to be driven cellular context (e.g., the *trans* environment) (**Fig. S3**). Evolutionarily, while RBPs are highly conserved, binding differences occur across species and are driven primarily by *cis* RNA evolution (**Fig. 5B**). These differences can emerge through evolution of primary motifs; however, substitutions within secondary binding sites can also lead to drastic binding differences across species (**Fig. 5E & Fig. S5D-J**).

RBP binding appears to be more complex than one might expect. While primary motifs can serve as “on/off” switches, the full mechanism is often more elaborate. For UNK specifically, the primary motif reported previously (UAGNNUUU) is only bound by UNK in 23% of all occurrences within the CDS, highlighting that other factors influence binding^27^. As has been reported before^53,54^, secondary motifs and local structure have evolved to modulate binding and regulation of individual transcripts. Given that many RBPs bind similar motifs, evolution of high-affinity binding sites presents an interesting balancing act. While enhanced accessibility and context may enhance individual RNA-RBP interactions, it may make these regions more accessible to other RBPs and thus enable RBP-RBP competition for binding.

While it’s difficult to look at the effects of single nucleotide variants (SNVs) on a global scale, the framework presented here should be broadly applicable for such studies. Indeed, previous work has shown that SNVs themselves can impact direct RBP-RNA interactions. When examining a “simple binding model,” SNVs within primary motifs can be understood to totally abrogate binding (**Fig. 6A**). When we look at a more “moderate binding model” and begin to consider secondary motif contributions and increased valency, we can see how UNK’s secondary motif may contribute to differential binding patterns across evolution (**Fig. 6B**). Finally, when examining the most “complex binding model,” which considers secondary structure as well, global structural rearrangements that affect motif access are the most striking examples (**Fig. 6C**). All three of these models — taking into account primary motifs, secondary motifs, and RNA structure — likely apply in different situations. Understanding selection for or against these complex features that impact binding will help explain how regulation is conserved or species-specific.

These binding differences have been observed throughout several previous studies, where RBP binding appears to be dynamic and cell-type specific^21,67–69^. Our findings support that genomic evolution of regulatory sites most frequently occurs in *cis*, rather than *trans*^6,7^, at least on shorter time frames (as shown with TFs^39^). However *cis* evolution can also result in *trans* evolution as regulatory elements like TFs and RBPs also have self-regulatory features^70^, and at longer distances *trans* variation may play a larger role. Additionally, the cellular environment is also changing. Thus, genomic evolution relies on a delicate balance of binding site mutations, protein conservation and expression, and cellular context.

UNK is a translational inhibitor that binds primarily in the CDS but can also bind in the 3’ UTR of its target mRNAs^27^. *In vitro*, UNK bound UTR sequences more tightly than CDS (**Fig. 2 & Fig. 3**), driven by the presence of more UNK motifs in UTRs (**Fig. S2A**). However, in cells, UNK binds primarily to CDS^27^; this preference may be driven by increased local concentration of UNK near CDS, resulting from its association with ribosomes^46^. UNK is not unique in exhibiting gene region preference^71^ and the mechanisms driving these preferences are not well understood. One recent study has demonstrated that UNK can associated with CCR4-NOT on some targets^47^, thus additional cellular factors like these may impact target selection.

The work presented here parallels studies performed on a handful of transcription factors^6,7,38,72–74^. For example, Schmidt and coworkers^6^ examined the binding profiles of two TFs in human, mouse, dog, opossum, and chicken and found that binding tends to be species-specific even when the proteins are highly conserved. Additionally, they observed that these species-specific binding preferences are largely due to *cis* sequence element changes across species^6^. For a more direct comparison, Odom *et al*.^7^ examined the binding profiles of four highly conserved TFs between human and mouse and found that while the TFs themselves are not readily changing, their DNA-interactome changes readily (up to 60%) often due to changes in motif content. Our work also highlights the complexity of translating defined regulatory elements from one species to another as the precise location of these sites may frequently change even when motifs appear to be conserved.

## Methods

### Expression and Purification of Recombinant UNK

Plasmid pGEX-GST-SBP-UNK (30-357)^26^ was transformed into Rosetta *E. coli* competent cells (Novagen). Cultures were grown to an OD of 0.8 in LB media, adjusted to 16°C, and induced with 0.5 mM IPTG (Thermo Scientific) for 24 hours. Cells were collected via centrifugation at 4,000 *x g* for 15 minutes and lysed in lysis buffer (200 mM NaCl, 5 mM DTT, 50 mM HEPES, 3 mM MgCl_2_, 2 mM PMSF, 1 Pierce™ protease inhibitor mini tablet/2 L; Thermo Scientific). The lysate was sonicated then incubated with 500 units/1 L culture Benzonase Nuclease (Sigma-Aldrich) for 30 minutes then with 5 units/1 L RQ1 RNase-free DNase (Promega) for 10 minutes at room temperature. NaCl concentration was adjusted to 1 M and the lysate was clarified by centrifugation at 17,800 *x g* for 30 minutes. 0.05% polyethyleneimine (PEI) was added to precipitate excess nucleotides and was centrifuged at 17,800 *x g* for 10 minutes.

Supernatant was passed over a 0.45-micron filter. Recombinant protein was purified via GST-trap FF column (GE). The column was washed in low salt buffer (300 mM NaCl, 50 mM HEPES), ATP buffer (300 mM NaCl, 50 mM HEPES, 5 mM ATP, 500 mM MgCl^2^), and high salt buffer (1 M NaCl, 50 mM HEPES). For 100vertRBNS only, 1:50 PreScission Protease (Cytiva) was loaded on column in cleavage buffer (20 mM HEPES, 100 mM NaCl, 5 mM DTT, 10% glycerol, 0.01% triton X-100) to cleave the GST-tag prior to SEC. Protein was incubated at 4°C overnight on-column. Cleaved SBP-tagged protein was eluted in cleavage buffer. For nsRBNS, proteins were eluted off column in glutathione buffer (50 mM Tris pH 8.0, 20 mM reduced glutathione, final pH = 7).

All proteins were concentrated via centrifugation to 500 μL before further purification via size exclusion chromatography [Superdex 200 Increase 10/300 GL (Cytiva)] in size exclusion buffer (20 mM HEPES, 1 M NaCl, 10 mM DTT, 0.01% triton X-100). 0.5-mL fractions were collected. Purity was assessed via SDS-PAGE (4-12% gradient) and Coomassie staining. Fractions corresponding to SBP-UNK (42.3 kDa) were pooled and dialyzed into 20 mM HEPES, 100 mM NaCl, and 5% glycerol. Concentration was determined via Pierce 660 nm assay (Thermo Scientific).

### iCLIP Data Analysis and Oligo Design

UNK iCLIP-seq data was obtained from E-MTAB-2279^27^. Mouse coordinates were converted from mm9 to mm10 and human coordinates were converted from hg19 to hg38 using liftOver^75^ (version 1.24.0) in RStudio^76^ (version 2023.03.0) with R platform^77^ (version 4.2.2). Peaks were selected such that only the maximum scoring peak within 20 nucleotides of other peaks would be recorded. This was done in a rolling fashion such that if several peaks were back-to-back, each within 20 nucleotides of each other, only the maximum scoring peak would be recorded. Peaks were mapped back to their respective genes/transcripts using RStudio package ‘AnnotationHub^78^’ (version 3.8.0) with ‘BSGenome.Hsapiens.NCBI.GRCh38^79^’ and ‘BSGenome.Mmusculus.UCSC.mm10^80^‘. Only peaks mapping to exons were included for subsequent analysis.

For overlap analysis, peaks were expanded to 101 nucleotides and sequences were obtained using ‘getSeq’ using BSgenomes mentioned above. TAG-containing regions were identified. Previously reported RNA-seq data for SH-SY5Y cells (Murn *et al*.:^27^ E-MATB-2277^81^) mouse brain tissue (ENCODE^60,82^: ENCSR000BZJ), and HeLa cells (ENCODE^60,82^: ENCSR552EGO) was mapped to the mm10 or hg38 genomes using STAR^83^ (version 2.7.10b) with default parameters. RSEM was used to calculate gene level expression values. Tximport was used to read RSEM output and transcripts per million (TPM) was used for comparison. Peaks were filtered to genes with >= 5 TPM in the respective cell lines. Gene level intersecting peaks (SH-SY5Y) were converted from hg38 to mm10 using the liftOver utility from UCSC^84^ then intersected with mouse iCLIP peaks using BEDtools^85^ (version 2.31.0). RStudio package ‘VennDiagram^86^’ was used to produce Venn diagrams.

For final oligo pool, sequences were expanded to 170 nucleotides and coding frame was determined where applicable. Using previously reported RBNS data for UNK^26^, sequences were recentered around the highest ranking *k*mer closest to the center. Final sequences were trimmed down to 120 nucleotides (**Fig. S1A**). A subset of oligos were selected from the bound and unbound human and mouse orthologs where the central UAG motifs were mutated to a CCG. Additionally, single and double chimeras were designed such that 10 (single) or 20 (double) nucleotides of the bound ortholog were placed into the unbound sequence. Due to the differences between human and mouse, the chimerization was not always perfect, meaning that placing 10 nt of human into the same exact syntenic mouse region may not be 100% accurate. This should be considered when interpreting the chimera data.

### Natural Sequence RNA Bind-n-Seq (nsRBNS)

Target RNAs were identified from iCLIP experiments performed in human and mouse neuronal cells^27^ (see above). An array of 24,254 natural sequence oligos was synthesized by Twist Biosciences and transcribed to RNA with T7 polymerase. RBNS was performed as previously described^26,52^. Briefly, MyOne Streptavidin T1 Dynabeads were washed in binding buffer (25 mM Tris pH 7.5, 150 mM KCl, 3 mM MgCl_2_, 0.01% tween, 500 μg/mL BSA, 1 mM DTT) and incubated with 0, 5, 50, or 500 nM recombinant GST-SBP-UNK. After 30-minute incubation, 1 μM RNA was added to the reaction. After 1 hour, UNK-RNA complexes were isolated and unbound RNA was washed away. Complexes were eluted in 4 mM biotin. The eluted RNA was reverse transcribed with Superscript III (Invitrogen) with RBNS RT primer (IDT; see below), amplified by PCR with Phusion DNA polymerase (NEB) with RBNS index primers and RBNS reverse primer I (see below), and sequenced on an Illumina HiSeq 300 instrument.

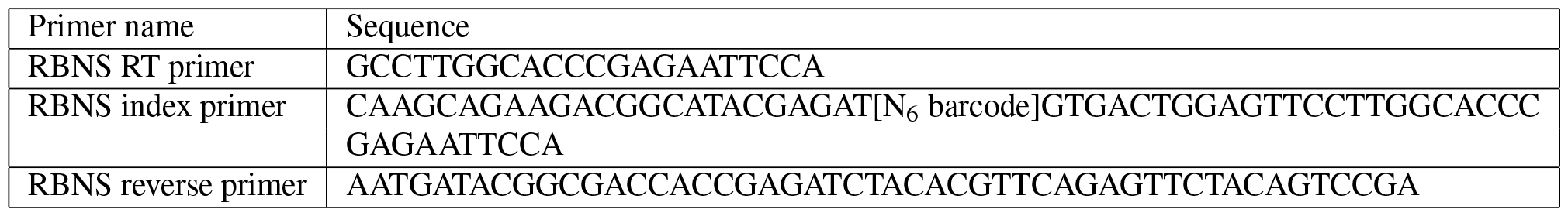

### nsRBNS Mapping and Enrichment Analysis

Reads were trimmed using fastx_toolkit^87^ (version 0.0.14) as needed. Mapping was performed with STAR^83^ (version 2.7.10b). STAR mapping parameters were set to –outFilterMultimapNMax 1 and –outFilterMismatchNmax 1 to generate counts files. Fasta file for reference was trimmed for adapters using seqtk^88^ (version 2.3.0) as needed. SAMtools^89^ (version 1.16) was used for processing alignment files as needed. Enrichment was calculated as frequency of an oligo in the protein bound sample divided by the frequency in the input.

### nsRBNS Data Analysis

Data was compiled and analyzed in RStudio^76^ (version 2023.03.0) with R platform^77^ (version 4.2.2). R packages ggplot2^90^ (version 3.4.1), ‘ggpattern^91^’ (version 1.0.1), ‘ggpubr^92^’ (version 0.6.0), and ‘ggrepel^93^’ (version 0.9.3) were used to make publication quality figures. Other RStudio packages — including ‘dplyr^94^’ (version1.1.0), ‘Hmisc^95^’ (version 4.8.0), ‘msa^96^’ (version 1.30.1), ‘org.Hs.eg.db^86^’ (version 3.17.0), ‘reshape2^97^’ (version 1.4.4), ‘rstatix^98^(Kassambara 2023b)’ (version 0.7.2), and ‘stringr^99^’ (version 1.4.4) — were used for data analysis as needed. GraphPad Prism (version 10) was also used to make publication quality figures as needed. Data tables used for all analyses with sequences, relevant iCLIP information, enrichment values, relevant sequence information, and relevant oligo information can be found in **Supplemental Table 1**.

### Ribosome Profiling Data Analysis

Ribosome profiling data was obtained from Shah *et al*^47^. Genes bound in both human and mouse, human only, or mouse only (human not bound) were identified via iCLIP as described above. For genes with multiple peaks, nsRBNS enrichment values were summed. Only human nsRBNS enrichments and Ribo-Seq log_2_ fold changes were used as ribosome profiling after UNK over-expression in mouse is not currently available.

### Mean Base Pair Probability Analysis

DNA sequences for all hg38 genes were obtained from Ensembl^100^. Genes not bound in human neuronal cells as identified by iCLIP^27^ were selected for subsequent analysis. For genes with multiple isoform sequences, only one was kept: This was done randomly with RStudio function “sample.” 120 nucleotide sequences were selected and centered around the downstream TAG motif just upstream of the stop codon to match the binding pattern of UNK which increases near the stop codon but does not bind the stop codon itself. Individual base pair probabilities were calculated with Vienna RNAfold^55^ –partfunc to calculate the partition function and base pairing probability matrix for both the CDS controls as well as “perfectly conserved,” “conserved,” “bound elsewhere,” and “not bound” sequences aligned perfectly at the central UAG. Mean base pair probability (bpp) was calculated for each category positionally and divided by CDS controls positionally to normalize. Mean bpp was further averaged across the central motif (UAG), five nucleotides upstream, and five nucleotides downstream.

### 100 Vertebrate DNA Pool Assembly

The top 250 human *in vivo* and *in vitro* bound sequences were selected from the nsRBNS experiment. UCSC’s BLAT^101^ was used to determine the chromosome as well as the start and stop position for each sequence. The start and stop positions were expanded out 65 nucleotides each such that the region was 250 nucleotides in total to account for insertions and deletions across species. Multiple alignment blocks were selected using maf_parse in PHAST module^102^ (version 1.5) against UCSC’s MAFs for 100 vertebrates. Sequences with less than 25 alignments were filtered out, and the remaining sequences were collapsed down with gaps removed. Sequences were aligned centrally in RStudio with package ‘msa’^103,104^ (version 1.30.1) and trimmed back to 120 nucleotides. RStudio packages ‘stringr’^99^ (version 1.5.0) and ‘data.table’^105^ (version 1.14.8) were also used for string manipulation and data table formation as needed. Finally, total motif mutants were included where all TAG motifs in human oligos were mutated to CCG to define a null cut-off for binding. From the starting 112 human oligos, we found 5,641 orthologous sequences.

### 100 Vertebrate RNA Bind-n-Seq

RBNS was performed similarly as described above and previously^26,52^. An array of 5,753 oligos corresponding to 112 human sequences and their 100 vertebrate alignments was ordered from Twist Biosciences. *In vitro* transcription was performed with T7 RiboMAX Express Large Scale RNA Production System (Promega) according to manufacturer protocols. RNA was purified via denaturing gel electrophoresis, eluted via RNA crush-n-soak into H_2_O, and concentrated with phenol chloroform extraction.

Binding reactions were assembled as follows: MyOne Streptavidin T1 Dynabeads (Thermo Fisher) were first washed in new RBNS binding buffer (25 mM Tris, pH 7.5, 150 mM KCl, 3 mM MgCl_2_, 0.01% triton-X 100, 500 μg/mL BSA, 20 units/mL SUPERase.In (Thermo Fisher)) then were incubated with 100 nM recombinant SBP-UNK for 30 minutes at 4°C. Following incubation, 1 μM RNA was added and incubated for 1 hour at 4°C. Unbound RNA was then washed away in new RBNS wash buffer (25 mM Tris pH 7.5, 150 mM KCl, 0.01% triton X-100, 20 units/mL SUPERase.In) three times. Bound RNA was eluted in 0.1% SDS and 0.3 mg/mL proteinase K (Thermo Fisher) at 60°C for 30 minutes. Elution was performed twice and the elutions were pooled. Following elution, reverse transcription was performed with Superscript IV (Invitrogen) with RBNS RT primer (see above) and amplified as described above. Sequencing was performed on an Illumina NextSeq 500.

### 100 Vertebrate nsRBNS Data Analysis

Data was analyzed similarly to nsRBNS data as discussed above with a few additions. RStudio package ‘ape’^106^ (version 5.7) was used to assemble 100 vertebrate phylogenic tree according to available data from UCSC^64,107^. Additionally, RStudio package ‘msa’^103,104^ (version 1.30.1) was used for sequence alignment and percent identity analysis. RStudio packages ‘ggmsa’^108^ (version 1.3.4), ‘ggprism^109^’ (version 1.0.4), and ‘scales^110^’ (version 1.2.1) were used to make publication quality alignment figures. Data tables used for all analyses with sequences, relevant species information, enrichment values, relevant sequence information, and relevant oligo information can be found in **Supplemental Table 2**. For protein percent similarity analysis, human protein sequences were pulled from UniProt^50^ and BLAST^48^ was used for all species alignments. The RBDs were annotated as amino acids 31 to 335 based on previous work by Murn *et al*^46^. All enrichments were normalized to their respective total motif mutant enrichment. Where indicated, delta enrichments were used for analysis and represent the divergence from the normalized human enrichment (log_2_[norm species/norm human]).

### eCLIP Peak Overlap Analysis

RNA-seq data for HepG2 and K562 normal cell lines were downloaded from ENCODE^60,82^ and mapped to hg38 with STAR^83^ (version 2.7.10b) using default parameters. Expression values were quantified using RSEM^111^ (version 1.3.1) and differential analysis was conducted using RStudio package ‘DESeq2’^111^ (version 1.40.1). Genes with no significant differences in expression between K562 and HepG2 cell lines were filtered using mean expression >10 and absolute log2 fold change (L2FC) <=1.

14 RBPs which have eCLIP peaks^61^ in HepG2 and K562 and an RBNS motif^26^ were selected. The peaks were downloaded from ENCODE (see table below) and filtered for enrichment of log_2_ fold change > 1 and p-value of <0.001. Each peak was extended by 50 base pairs upstream to account for experimental limitations of eCLIP^61^. Replicates for each cell line were combined and filtered for the ones that fell within the genes that did not have significantly differential expression between HepG2 and K562. BEDtools^85^ (version 2.31.0) was used to collapse the overlapping peaks that were within 20 base pairs into a single peak spanning the region. Sequences under the combined peaks were taken from RStudio package ‘BSgenome.Hsapiens.NCBI.GRCh38’^79^ (version 1.3.1). Peaks were grouped into two groups based on presence or absence of RBNS motif within the peak^26^.

Peaks were further grouped based on whether they overlapped exons or not. Exon annotations for hg38 (v109) genome were obtained from RStudio package ‘Annotation Hub’^78^ (version 3.8.0) and a peak was assigned as an exonic peak if it had at least 20 base pair overlap with an exon. Thus, peaks for an RBP in each cell line were grouped into four following groups: peaks within an exon and had an RBNS motif, peaks within an exon and did not have an RBNS motif, peaks not within an exon and had an RBNS motif, and peaks not within an exon and did not have an RBNS motif. For each of these groups, we calculated the number of overlapping peaks between K562 and HepG2. Since the proportion of overlapping peaks between two cell lines is relative to total peaks identified in each cell line, we took the maximum after calculating the proportion using both cell lines respectively. Additionally, similar analysis was conducted for overlaps between parent genes of the peaks as well (**Fig. S3F**).

| RNA-Seq | Cell Line | Rep |
| --- | --- | --- |
| ENCFF267RKD,ENCFF455VYN | K562 | 1 |
| ENCFF606ZTR,ENCFF444KCV | K562 | 2 |
| ENCFF713MNU,ENCFF478DZZ | HepG2 | 1 |
| ENCFF936SLY,ENCFF446UEC | HepG2 | 2 |

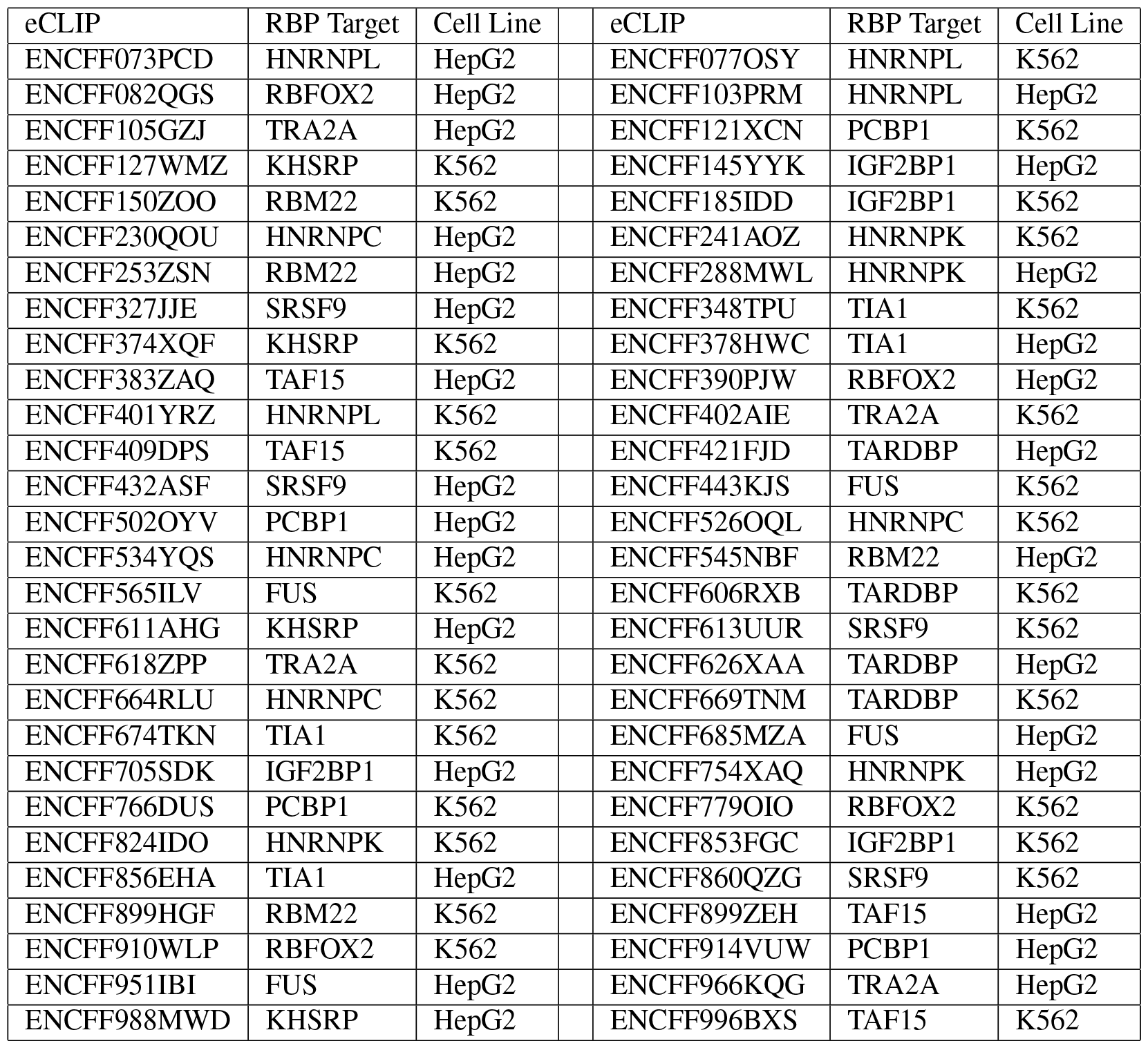

| eCLIP | RBP Target | Cell Line | eCLIP | RBP Target | Cell Line |
| --- | --- | --- | --- | --- | --- |
| ENCFF073PCD | HNRNPL | HepG2 | ENCFF077OSY | HNRNPL | K562 |
| ENCFF082QGS | RBFOX2 | HepG2 | ENCFF103PRM | HNRNPL | HepG2 |
| ENCFF105GZJ | TRA2A | HepG2 | ENCFF121XCN | PCBP1 | K562 |
| ENCFF127WMZ | KHSRP | K562 | ENCFF145YYK | IGF2BP1 | HepG2 |
| ENCFF150ZOO | RBM22 | K562 | ENCFF185IDD | IGF2BP1 | K562 |
| ENCFF230QOU | HNRNPC | HepG2 | ENCFF241AOZ | HNRNPK | K562 |
| ENCFF253ZSN | RBM22 | HepG2 | ENCFF288MWL | HNRNPK | HepG2 |
| ENCFF327JJE | SRSF9 | HepG2 | ENCFF348TPU | TIA1 | K562 |
| ENCFF374XQF | KHSRP | K562 | ENCFF378HWC | TIA1 | HepG2 |
| ENCFF383ZAQ | TAF15 | HepG2 | ENCFF390PJW | RBFOX2 | HepG2 |
| ENCFF401YRZ | HNRNPL | K562 | ENCFF402AIE | TRA2A | K562 |
| ENCFF409DPS | TAF15 | K562 | ENCFF421FJD | TARDBP | HepG2 |
| ENCFF432ASF | SRSF9 | HepG2 | ENCFF443KJS | FUS | K562 |
| ENCFF502OYV | PCBP1 | HepG2 | ENCFF526OQL | HNRNPC | K562 |
| ENCFF534YQS | HNRNPC | HepG2 | ENCFF545NBF | RBM22 | HepG2 |
| ENCFF565ILV | FUS | K562 | ENCFF606RXB | TARDBP | K562 |
| ENCFF611AHG | KHSRP | HepG2 | ENCFF613UUR | SRSF9 | K562 |
| ENCFF618ZPP | TRA2A | K562 | ENCFF626XAA | TARDBP | HepG2 |
| ENCFF664RLU | HNRNPC | K562 | ENCFF669TNM | TARDBP | K562 |
| ENCFF674TKN | TIA1 | K562 | ENCFF685MZA | FUS | HepG2 |
| ENCFF705SDK | IGF2BP1 | HepG2 | ENCFF754XAQ | HNRNPK | HepG2 |
| ENCFF766DUS | PCBP1 | K562 | ENCFF779OIO | RBFOX2 | K562 |
| ENCFF824IDO | HNRNPK | K562 | ENCFF853FGC | IGF2BP1 | K562 |
| ENCFF856EHA | TIA1 | HepG2 | ENCFF860QZG | SRSF9 | K562 |
| ENCFF899HGF | RBM22 | K562 | ENCFF899ZEH | TAF15 | HepG2 |
| ENCFF910WLP | RBFOX2 | K562 | ENCFF914VUW | PCBP1 | HepG2 |
| ENCFF951IBI | FUS | HepG2 | ENCFF966KQG | TRA2A | HepG2 |
| ENCFF988MWD | KHSRP | HepG2 | ENCFF996BXS | TAF15 | K562 |

### Fluorescence Polarization (FP)

RNA oligos were synthesized by Integrated DNA Technologies (IDT) with a 3’ 6-FAM label (see below) and incubated at 5 nM with serially diluted recombinant SBP-UNK (10.9, 50.8, 152, 457 pM, 1.37, 4.12, 12.3 37.0, 111, 333, or 1000 nM) in FP binding buffer (20 mM HEPES, 5 mM DTT, 137.5 mM NaCl, 0.01% triton X-100, 10 ng/μL BSA, 2 units/mL SUPERase•In™; Thermo Scientific) for 15 min at 4°C. Plates were centrifuged at 1000 *x g* for 1 min and fluorescence polarization was measured with a PHERAstar plate reader (BMG Labtech) at 25°C. Data were fit to a single site binding model and a K_D_ was determined.

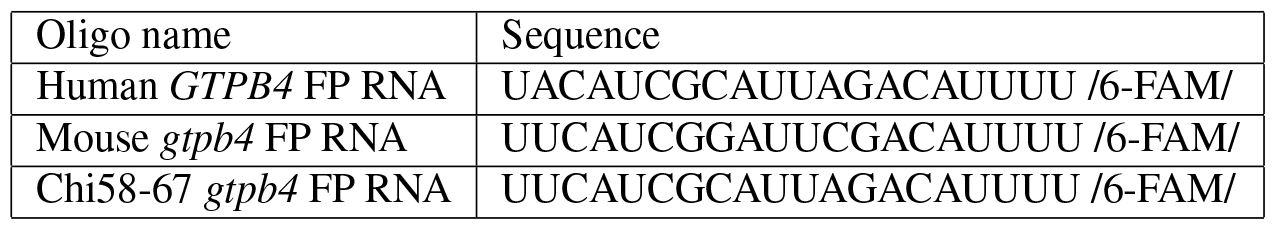

### *In vitro* Transcription of RNA for qPCR Binding Assay

DNA fragments for wild-type and mutant GART were ordered from IDT (see below) and PCR amplified with Phusion DNA polymerase (New England Biolabs) with FWD_adapter and REV_adapter primers from IDT (see below), resulting in full-length DNA oligos for *in vitro* transcription. DNA was purified via agarose gel extraction, and transcribed with a T7 RiboMAX Express Large Scale RNA Production System (Promega). RNA was purified with RNA crush-n-soak and concentrated with phenol chloroform isolation.

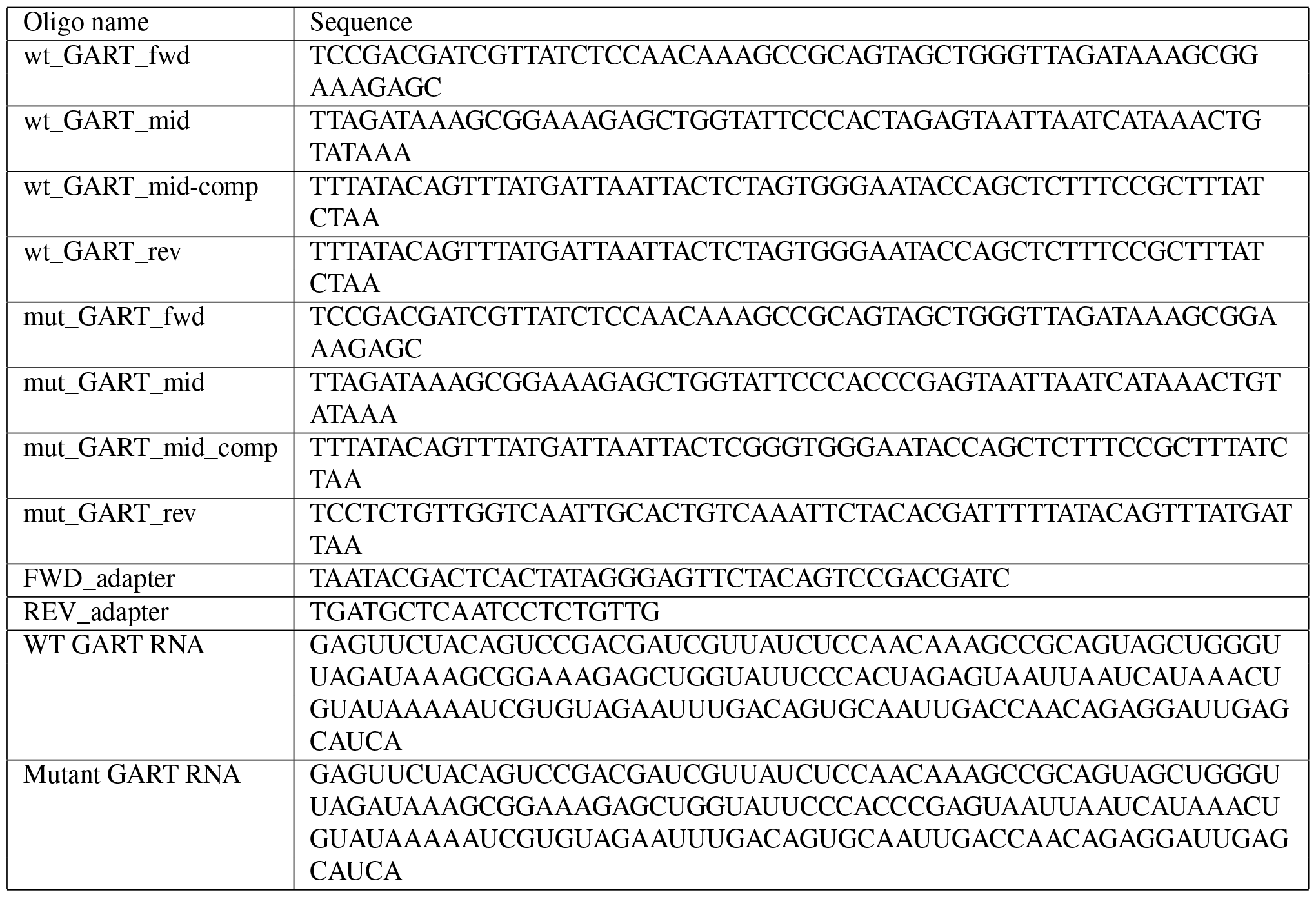

### qPCR Binding Assay

We performed our qPCR-based binding assay as previously described^112^ with a few modifications. Dynabeads MyOne Streptavidin T1 magnetic beads (Thermo Fisher) were washed in blocking buffer (25 mM Tris pH 8.0, 150 mM KCl, 3 mM MgCl_2_, 1 mg/mL BSA, 2 units/1 μL SUPERase-In (Invitrogen), and 1 mg/mL yeast tRNA (Fisher Scientific)), and then in qPCR binding buffer (25 mM Tris pH 8.0, 150 mM KCl, 3 mM MgCl_2_, 1 mg/mL BSA, 2 units/1 mL SUPERase-In, and 50 nM random sequence RNA). Beads were incubated with two concentrations of SBP-UNK (167 and 1500 nM) at 25°C for 10 minutes. Bead-protein complexes were separated on the magnet and resuspended in 0.1 nM RNA. Beads, protein, and RNA were incubated at 25°C for 30 minutes. Unbound RNA was removed, and bound RNA was eluted in 4 mM biotin and 25 mM Tris, pH 8.0 at 37°C for 30 minutes. Reverse transcription was performed on unbound and bound RNA with iScript Reverse Transcription Supermix (Bio-Rad) following manufacturer’s protocols with qPCR_REV primer (see below). An RNA calibration curve was assembled at RT with the following amounts of RNA: 45.7 fM, 0.137, 0.412, 1.23, 3.70, 11.1, 33.3, and 100 pM. RT reactions were diluted 2-fold and qPCR was performed in duplicate with SsoAdvanced SYBR Green Supermix (Bio-Rad) according to manufacturer’s protocols with qPCR_FWD and qPCR_REV primers from IDT (see below). The threshold cycle (C_t_) was determined using Bio-Rad’s CFX Maestro software, and fraction bound was determined against the RNA calibration curve.

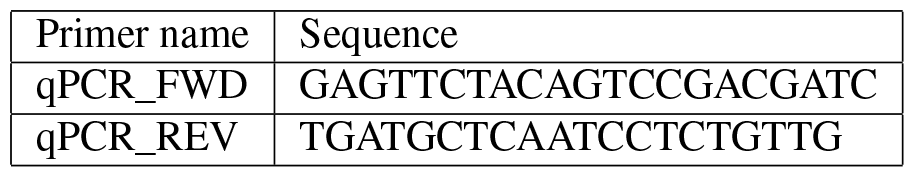

## Acknowledgements

We would like to thank Michael Love, Peter Sudmant, Matthew Taliaferro, and Eric Van Nostrand for helpful and insightful feedback on this manuscript. This work was in part supported by NIH T32 GM008570 (S.E.H.) and R35GM142864 (D.D.) as well as startup funds from UNC Chapel Hill to (D.D.).

## Author Contributions

Conceptualization, S.E.H., M.S.A., C.B., and D.D.; Methodology, S.E.H., M.S.A., M.M.A., and D.D.; Software, S.E.H., M.S.A., G.G., F.F.C., and D.D.; Validation, S.E.H., M.S.A., G.G., M.M.A., and D.D.; Formal Analysis, S.E.H., M.S.A., F.F.C., and D.D.; Investigation, S.E.H., M.S.A., M.M.A., and D.D.; Resources, C.B. and D.D.; Data Curation, S.E.H., M.S.A., G.G., J.M., and D.D.; Writing – Original Draft, S.E.H. and D.D.; Writing – Review & Editing, S.E.H., M.S.A., G.G., J.M., M.M.A., C.B.B., and D.D.; Visualization, S.E.H., G.G., and D.D.; Supervision, C.B.B. and D.D.; Funding Acquisition, D.D.

