## Supplemental for "Understanding species-specific and conserved RNA-protein interactions *in vivo* and *in vitro*"

### Index of Supplementary Information for Harris *et al*.

Supplementary Notes

Supplementary References

Supplementary Figures

- Supp. Figure 1 → relates to Figure 1: Design and validation of natural sequence RNA Bind-n-Seq
- Supp. Figure 2 → relates to Figure 2: Analysis of species-specific binding patterns
- Supp. Figure 3 → relates to Figure 3: Analysis of species-specific syntenic motif-level binding patterns
- Supp. Figure 4 → relates to Figure 4: Analysis of regional impacts on binding
- Supp. Figure 5 → relates to Figure 5: Evolutionary conservation of binding

Supplementary Download Items

- Supp. Table 1 → relates to Figure 1-4: nsRBNS data table used for all analyses with sequences, relevant iCLIP information, enrichment values, zscores, relevant sequence information, and relevant oligo information
- Supp. Table 2 → relates to Figure 5: 100vert RBNS data table used for all analyses with sequences, relevant species information, enrichment values, relevant sequence information, and relevant oligo information

#### SUPPLEMENTARY NOTES

##### Supp. Note 1

In our nsRBNS experiments, we have observed that human oligos are better bound than mouse oligos on a global scale. This can be observed for all oligos, bound alone, or unbound alone. The precise nature of these differences is difficult to define, however a few factors may contribute. While we can't be certain, one likely explanation might be in a technical challenge of performing iCLIP from mouse brain tissue vs human neuronal cell lines. It is plausible that the identified mouse binding sites were more challenging to derive due to tissue processing and crosslinking efficiencies as well as the abundance of UNK itself. These differences are apparent in some of our comparisons and limited our ability to test the impact of chimerizing mouse bound sequences into human not bound sequences to test if any enhanced UNK binding. Within these analyses, we found that while some regions from mouse bound sequences could enhance human binding, specific positions were not as impactful as we observed when chimerizing human into mouse (**Fig. 4B,C**). Generally, trends of binding patterns were overall weaker for mouse binding sites.

Another potential mechanism driving this is effect are inherent protein differences: Human recombinant protein was used for all *in vitro* experiments; however iCLIP was performed *in vivo* with species-specific proteins[1]. At the amino acid sequence level, human and mouse UNK are 95% identical and 96% similar overall while the RBDs are 99% identical[2,3]. Within the RBDs, only one amino acid is non-similar (Q321 in human, P321 in mouse). Structurally, this amino acid does not lie on the RNA-binding surface and likely does not directly contribute to RNA-protein interactions[4]. However, this single amino acid difference may affect global structural orientation, and therefore indirectly alter RNA-protein interactions *in vivo*. Indeed, previous work on transcription factors has demonstrated preferential binding of human sequences by human protein and vice versa[5].

Cellular context differences: Individual RNA and protein concentrations vary across cell-types and species. Additionally, species-specific alternative splicing can result in sequence differences and isoform expression level changes across cell-types and species and have been largely correlated with genomic evolution[6–9]. These may be particularly important when comparing tissues to cell lines. Cell type mRNA composition and heterogeneity is likely an important consideration when assessing iCLIP from tissue samples. It is plausible that cell-type specific binding events within a tissue dampen the degree to which binding to any one site is detected. Previous work has demonstrated that RNA-protein interactions across tissue culture cells versus isolated tissue can vary drastically, even when similar cell types are considered[10].

Since nsRBNS does not take into consideration any competitory binding, cofactors, salt concentrations, diffusion differences, etc. between cells and purely measures RNA-protein interactions, these factors may be different across species, even when similar cell types are examined.

**Supp. Note 2**

In our 100vert RBNS experiment, we observe a decreased range of enrichments (-3 to 3 on $\log_2$ scale) versus nsRBNS (-6 to 6 on $\log_2$ scale). RBNS is inherently a zero-sum experiment (aka gain must equal loss). While both nsRBNS and 100vert RBNS are impacted by the nature of these experiments, oligo design for 100vert RBNS led to a greater impact (*i.e.*, dampening of signal).

For comparison, in nsRBNS only ~5,000 (20%) out of ~25,000 oligos were predicted to bind, whereas ~7,000 (28%) had unknown binding capabilities, while the remaining ~13,000 (52%) were expected to be non-binders. While non-binders were primarily included as controls and for validation purposes, this allowed for less competition within the oligo pool, and therefore a larger separation of enrichment values.

In contrast, only ~10% of expected non-binders were included for 100vert RBNS. Further, 100vert RBNS was designed from the top bound oligos in nsRBNS, further narrowing expected binding differences across oligos. This prevalence of high-capacity binders led to an earlier saturation of UNK protein with high competition potential between the oligos. The increased competition dampened the overall signal of the experiment, resulting in a decreased range of enrichments measured.

### Supplemental Figure 1

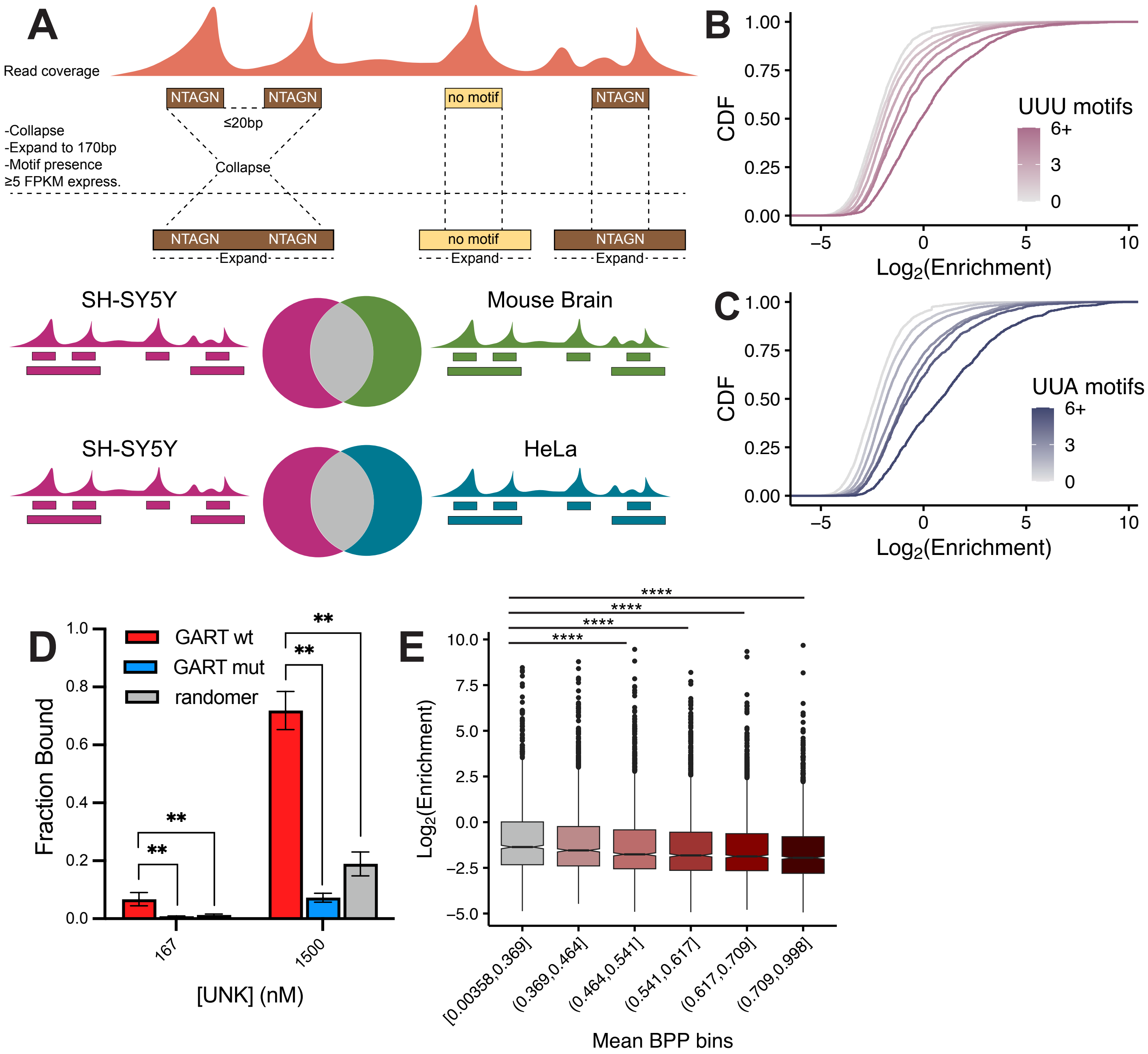

### Supplemental Figure 2

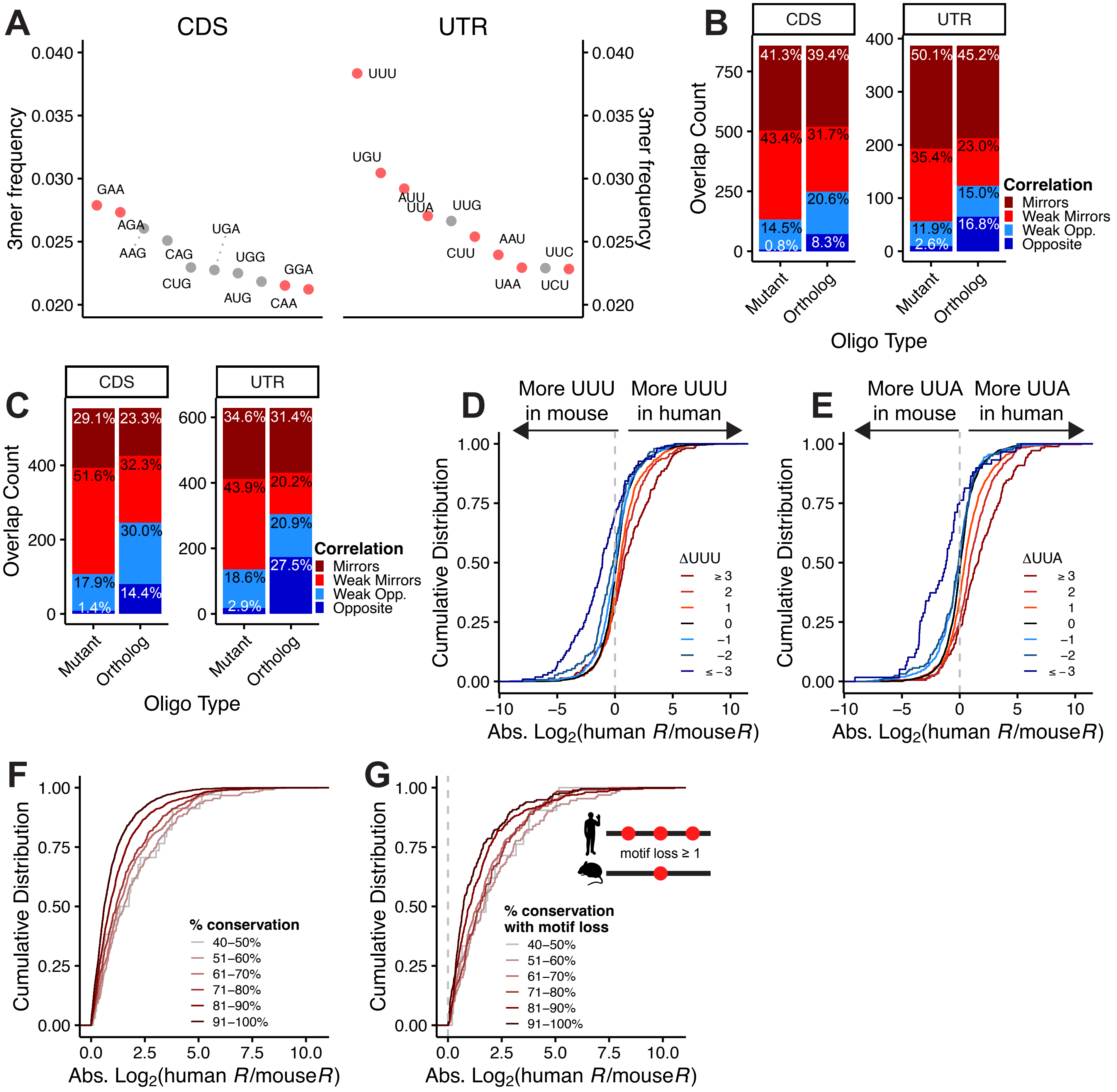

### Supplemental Figure 3

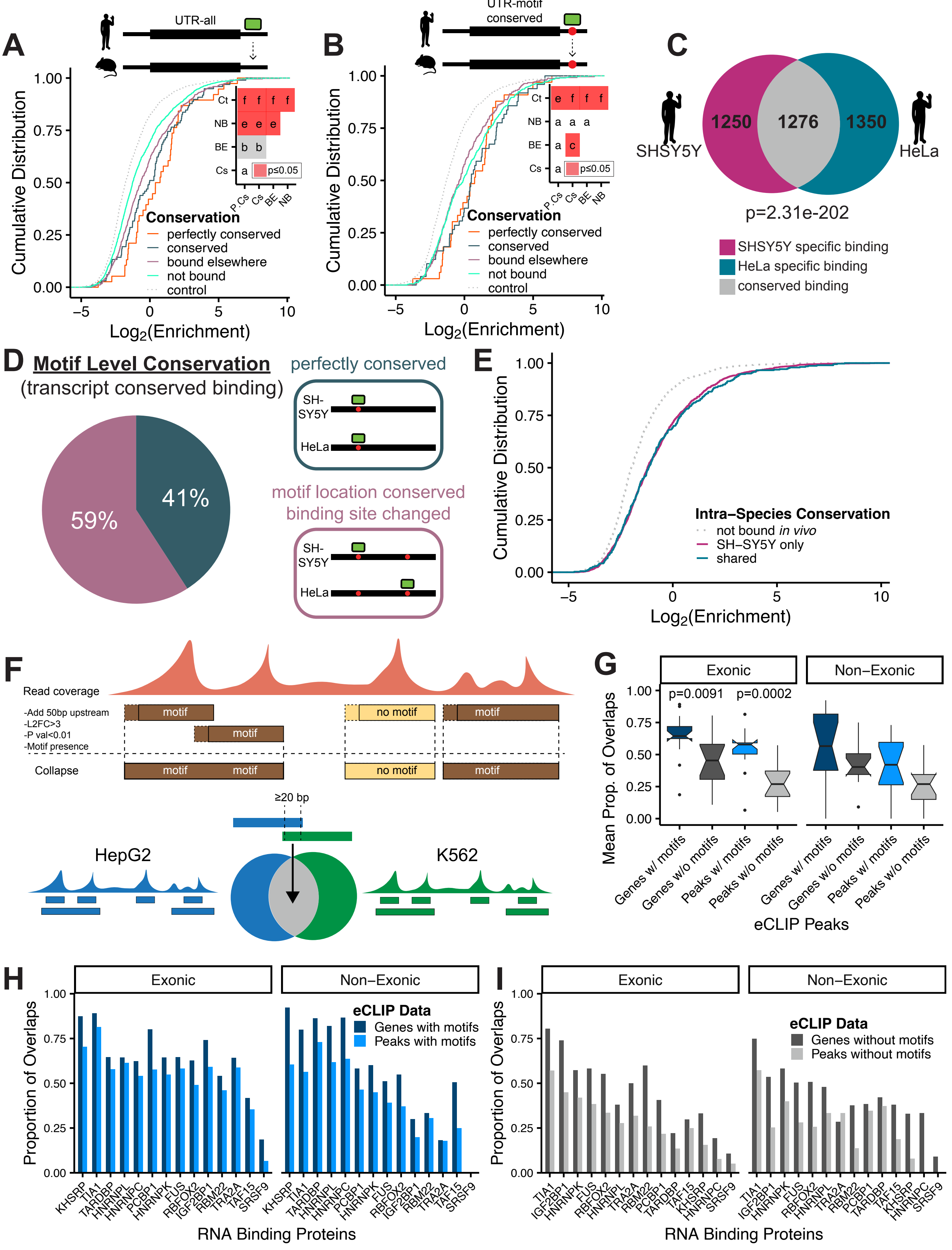

### Supplemental Figure 4

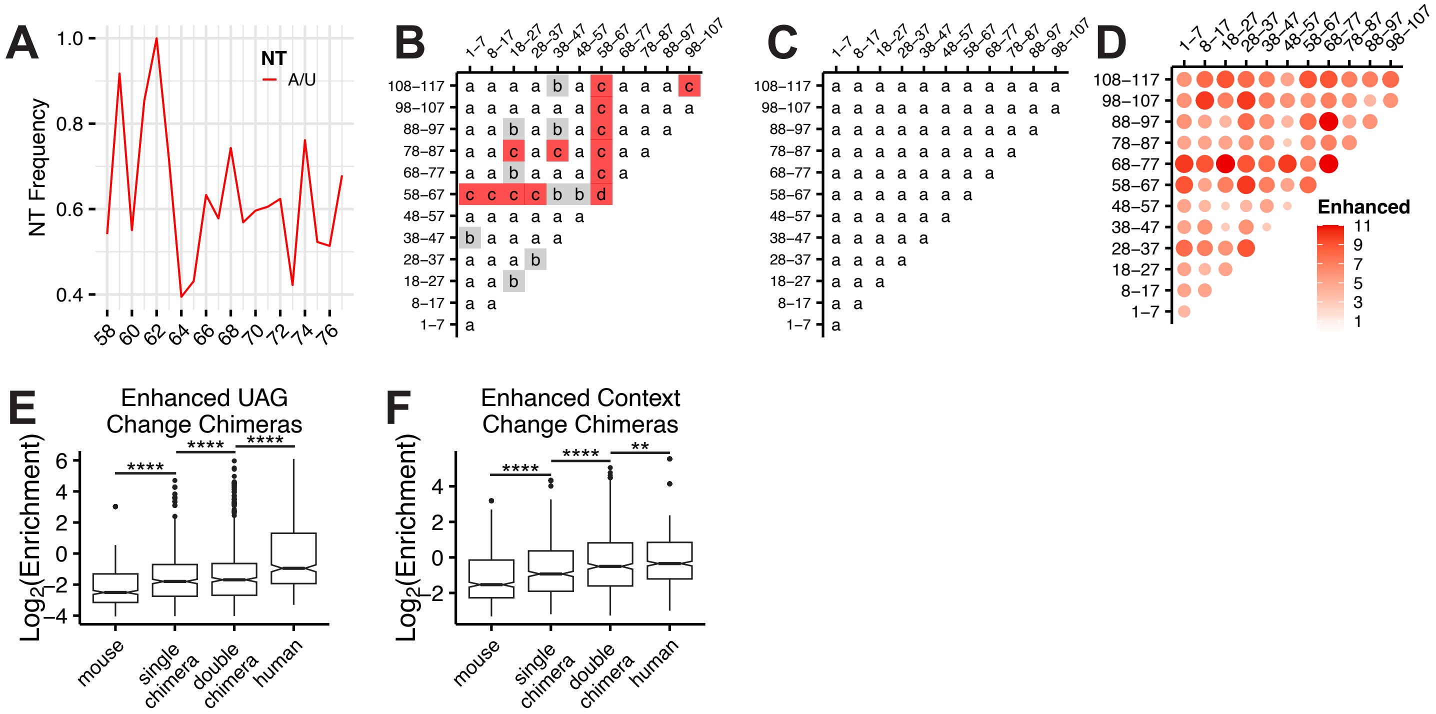

### Supplemental Figure 5

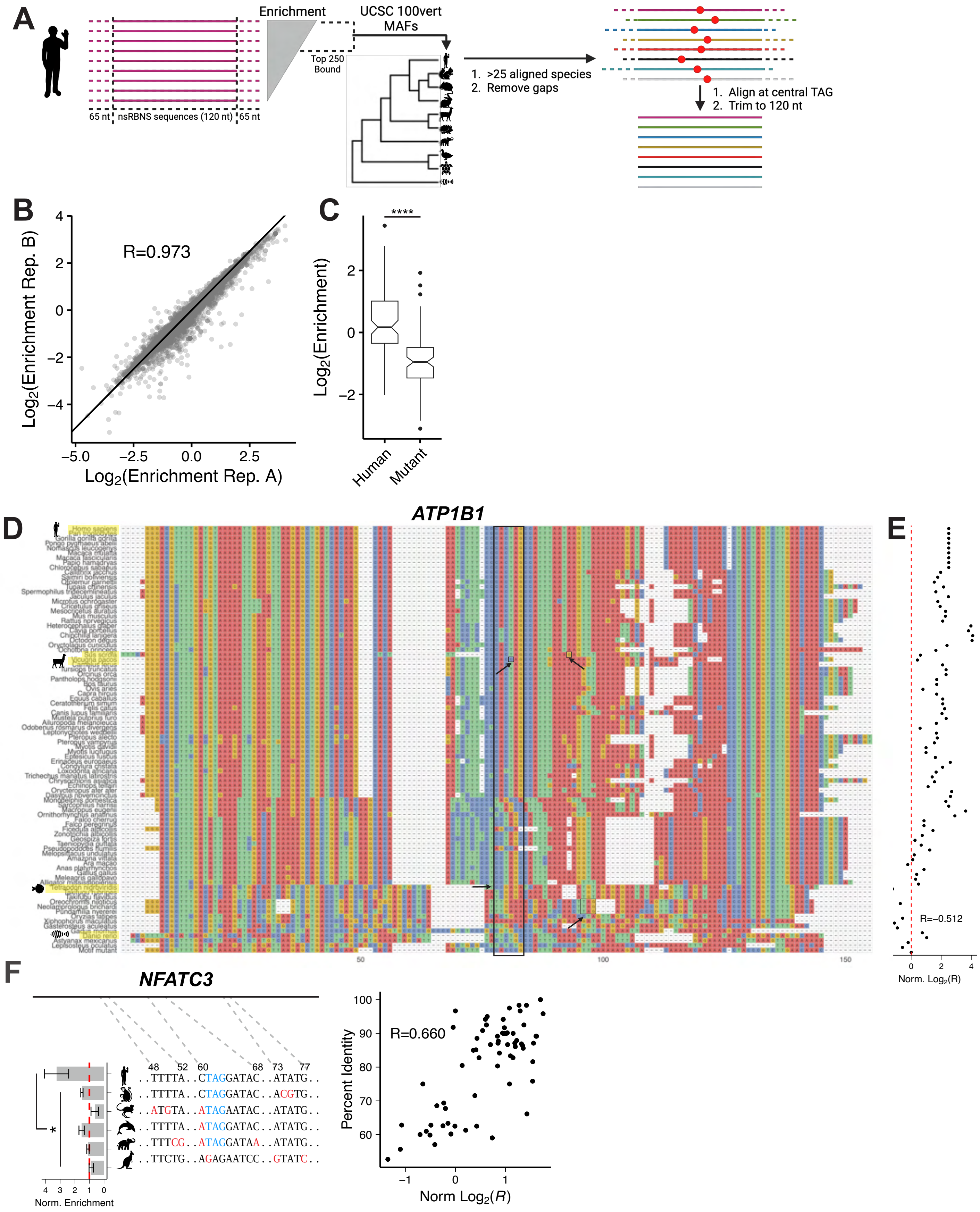

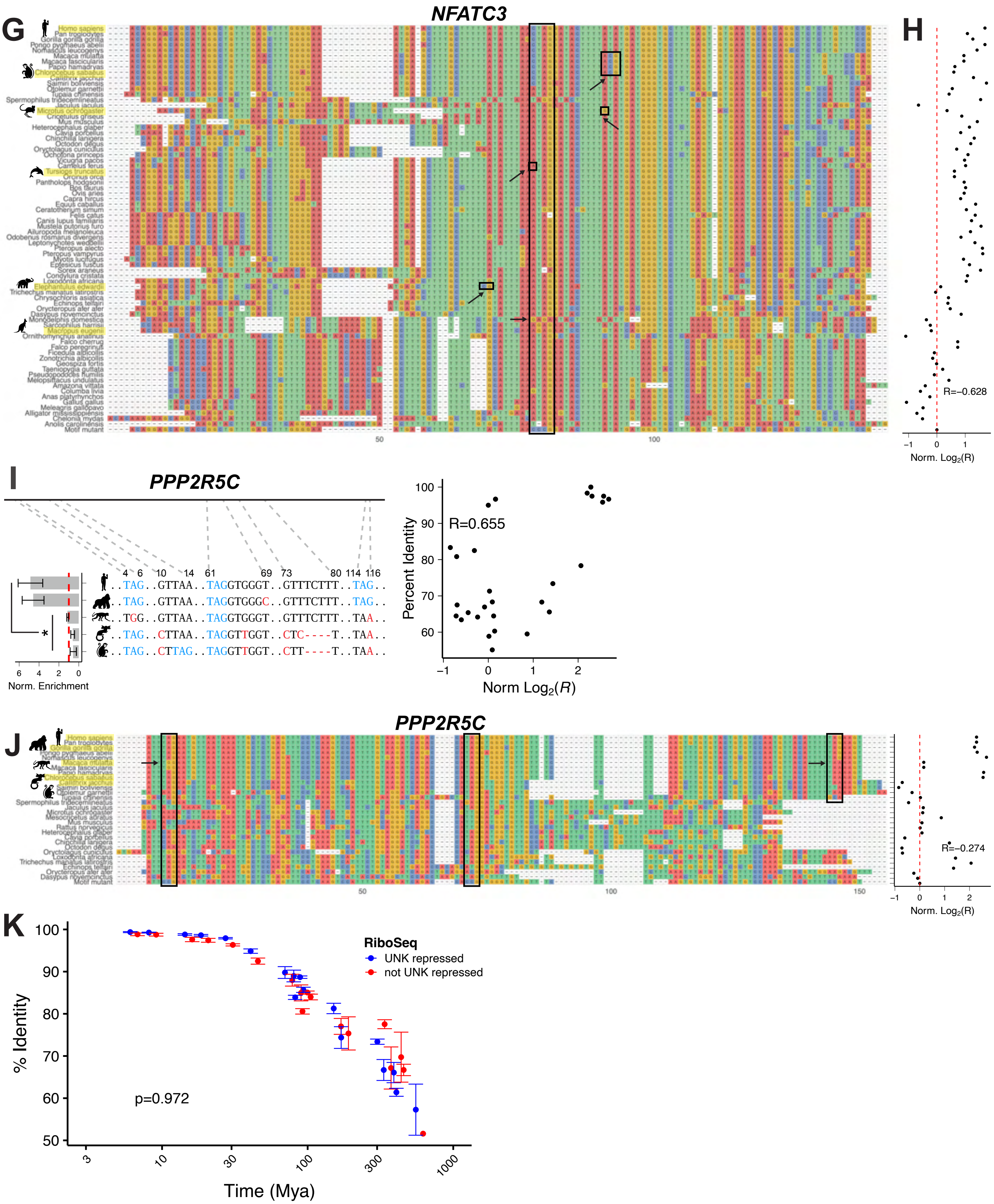
G
NFATC3
H
R=−0.628
Norm. Log2(R)
I
PPP2R5C
4 6 10 14 61 69 73 80 114 116
..TAG..GTTAA..TAGGTGGGT..GTTTCTTT..TAG..
..TAG..GTTAA..TAGGTGGGC..GTTTCTTT..TAG..
..TGG..GTTAA..TAGGTGGGT..GTTTCTTT..TAA..
..TAG..CTTAA..TAGGTTGGT..CTC----T..TAA..
..TAG..CTTAG..TAGGTTGGT..CTT----T..TAA..
Norm. Enrichment
R=0.655
Percent Identity
Norm Log2(R)
J
PPP2R5C
R=−0.274
Norm. Log2(R)
K
RiboSeq
UNK repressed
not UNK repressed
p=0.972
% Identity
Time (Mya)

#### SUPPLEMENTARY LEGENDS

**Supp. Fig. 1. Design and validation of natural sequence RNA Bind-n-Seq.** A) Diagram of iCLIP peak overlap analysis method between human neuronal cells (SH-SY5Y) and mouse brain tissue or human HeLa cells. B-C) Cumulative distribution function of $\log_2$ enrichment of all oligos separated by B) UUU and C) UUA motif content. D) *in vitro* qPCR fraction bound for wild type and mutant *GART* RNA oligos incubated with UNK at 167 and 1500 nM. Significance was determined via one-sided Wilcox test ($p \leq 0.01$). E) Box and whisker plot of $\log_2$ enrichment of all oligos separated by quantile-binned mean base pair probability (BPP) of the central region (54-64). Significance determined via KS test and corrected for multiple corrections via the BH procedure. All comparisons to lowest bin are significant ($p \leq 0.0001$).

**Supp. Fig. 2. Analysis of species-specific binding patterns.** A) Scatter plot of the *k*mer frequency of the top ten 3mers of all (left) CDS and (right) UTR oligos colored by UNK bound *k*mer as identified via RBNS[11]. B-C) Bar plot of *in vivo* binding versus *in vitro* binding patterns for "motif mutant" and "orthologous" oligos versus B) human and C) mouse *in vivo* bound oligos. "Mirrors" correlation defined as ≥ 2-fold change of *in vivo* bound over *in vivo* unbound, "weak mirrors" defined as < 2 and ≥ 1 fold change, "weak opposite" defined as < 1 and > 0.5 fold change, and "opposite" defined as ≤ 0.5 fold change. D-E) Cumulative distribution function of $\log_2$ fold enrichment change of *in vivo* bound over *in vivo* not bound oligos separated by D) ∆UUU and E) ∆UUA content. F-G) Cumulative distribution function of $\log_2$ fold enrichment change of *in vivo* bound over *in vivo* not bound oligos separated by percent conservation for F) all and G) *k*mer loss cross-species comparisons.

**Supp. Fig 3. Analysis of species-specific syntenic motif level binding patterns.** A-B) Cumulative distribution function of $\log_2$ enrichment of control (light grey; dotted), not bound (teal), bound elsewhere (purple), conserved (blue), and perfectly conserved (orange) for A) all UTR and

B) motif conserved UTR oligos. Insets show significance values for all comparisons via KS test and corrected for multiple comparisons via the BH procedure. Red denotes significant ($p \leq 0.05$) and gray denotes nearing significant ($p \leq 0.1$). Values are as follows: a (ns), b ($p \leq 0.1$), c ($p \leq 0.05$), e ($p \leq 0.001$), f ($p \leq 0.0001$). C) Transcript level conservation of iCLIP UNK hits between human neuronal cells (SH-SY5Y) and human epithelial cells (HeLa). Significance determined via hypergeometric test. D) Motif level conservation of iCLIP UNK hits between human neuronal cells (SH-SY5Y) and human epithelial cells (HeLa). E) Cumulative distribution function of $\log_2$ enrichment of human not bound *in vivo* (light grey; dotted), SH-SY5Y-specific oligos (purple), and SH-SY5Y and HeLa shared oligos (bliue) F) Diagram of eCLIP peak overlap analysis between HepG2 and K562 data. G) Mean proportion of overlaps of bound eCLIP genes and peaks between available HepG2 and K562 data[12] for 14 RBPs with and without known motifs from RBNS[11]. Data split into exonic and non-exonic peaks (see **Methods**). H) Overlap of bound eCLIP genes and peaks between available HepG2 and K562 data[12] for 14 RBPs with known motifs from RBNS[11]. Data split into exonic and non-exonic peaks (see **Methods**). I) Overlap of bound eCLIP genes and peaks between available HepG2 and K562 data[12] for 14 RBPs without known motifs from RBNS[11]. Data split into exonic and non-exonic peaks (see **Methods**).

**Supp. Fig. 4. Analysis of regional impacts on binding.** A) A/U nucleotide frequency of the central region (pos58-77) of enriched “Context Change” chimeras. B-C) Heat map of significance for all single and double B) “UAG Change” and C) “Context Change” chimeras with significant binding changes ($p \leq 0.05$, red) and nearing significant ($p \leq 0.1$, grey). Values are as follows: a (ns), b ($p \leq 0.1$), c ($p \leq 0.05$), d ($p \leq 0.01$), e ($p \leq 0.001$), f ($p \leq 0.0001$). Significance was determined via paired, one-sided Wilcox test and corrected for multiple comparisons via the BH procedure. D) Heat map of number of “Context Change” double chimeras enhanced upon chimerization. E-F) Box and whisker plot of $\log_2$ enrichment of mouse, single chimera, double chimera, and human oligos where both the single and double chimeras showed improved binding over mouse for E)

"UAG Change" and F) "Context Change" chimeras. Significance was determined via paired, one-sample, Wilcox test. Statistical marks are as follows: ** — $p \leq 0.01$, **** — $p \leq 0.0001$.

**Supp. Fig. 5. Evolutionary Conservation of Binding.** A) Design of 100 vertebrate DNA pool. B) Correlation plot of two experimental 100 vertebrate nsRBNS replicates. Pearson's correlation coefficient included. C) Box and whisker plot of $\log_2$ enrichment for human and total motif mutants. Significance was determined via paired, one-sided Wilcox test. Statistical marks are as follows: **** — $p \leq 0.0001$. D) Full multiple sequence alignment for *ATP1B1*. Related to figure 5E. E) Normalized $\log_2$ enrichment of *ATP1B1*. F) (left) Multiple sequence alignment for *NFATC3 for Homo sapiens, Chlorocebus sabaeus, Jaculus jaculus, Tursiops truncatus, Elephantulus edwardii*, *and Macropus eugenii* with normalized enrichment by species. (right) Percent RNA sequence identity (Y-axis) versus normalized delta $\log_2$ enrichment (X-axis). Pearson's correlation coefficient included. G) Full multiple sequence alignment for *NFATC3*. H) Normalized $\log_2$ enrichment of *NFATC3*. I) (left) Multiple sequence alignment for *PPP2R5C* for *Homo sapiens, Gorilla gorilla gorilla, Macaca mulatta, Callithrix jacchus,* and *Saimiri boliviensis* with normalized enrichment by species. (right) Percent RNA sequence identity (Y-axis) versus normalized delta $\log_2$ enrichment (X-axis). Pearson's correlation coefficient included. J) (left) Multiple sequence alignment for *PPP2R5C*. (right) Normalized $\log_2$ enrichment of *PPP2R5C*. F) Plot of percent identity by evolutionary distance, X-axis plotted on $\log_{10}$ scale. Error bars show SEM. Data was separated by regulation as determined via RiboSeq where blue is higher than average $\log_2$ fold change (>-0.9) and red is less than average $\log_2$ fold change (<-0.9). Significance was determined via KS test.
